# Signal recovery in single cell batch integration

**DOI:** 10.1101/2023.05.05.539614

**Authors:** Zhaojun Zhang, Divij Mathew, Tristan Lim, Kaishu Mason, Clara Morral Martinez, Sijia Huang, E. John Wherry, Katalin Susztak, Andy J. Minn, Zongming Ma, Nancy R. Zhang

## Abstract

Data integration to align cells across batches has become a cornerstone of single cell data analysis, critically affecting downstream results. Yet, how much biological signal is erased during integration? Currently, there are no guidelines for when the biological differences between samples are separable from batch effects, and thus, data integration usually involve a lot of guesswork: Cells across batches should be aligned to be “appropriately” mixed, while preserving “main cell type clusters”. We show evidence that current paradigms for single cell data integration are unnecessarily aggressive, removing biologically meaningful variation. To remedy this, we present a novel statistical model and computationally scalable algorithm, CellANOVA, to recover biological signal that is lost during single cell data integration. CellANOVA utilizes a “pool-of-controls” design concept, applicable across diverse settings, to separate unwanted variation from biological variation of interest. When applied with existing integration methods, CellANOVA allows the recovery of subtle biological signals and corrects, to a large extent, the data distortion introduced by integration. Further, CellANOVA explicitly estimates cell- and gene-specific batch effect terms which can be used to identify the cell types and pathways exhibiting the largest batch variations, providing clarity as to which biological signals can be recovered. These concepts are illustrated on studies of diverse designs, where the biological signals that are recovered by CellANOVA are shown to be validated by orthogonal assays. In particular, we show that CellANOVA is effective in the challenging case of single-cell and single-nuclei data integration, where the recovered biological signals are replicated in an independent study.

## Introduction

Over the last decade, single cell experiments have become routine in the biomedical field. Early efforts in single cell profiling focused on atlas building: Samples from one or a few replicates of a biological system are taken, with the goal of comprehensively mapping the cell types that make up the system. While such efforts continue, standardization and commercialization of single cell technologies have enabled large-cohort, population-scale studies to interrogate the cell types and cell type-level changes that underpin diseases.

Batch effects (also called “unwanted variation”) are pervasive in single cell studies, and the integration of cells across samples to remove batch effects is a critical step in any analysis pipeline (1–3). Although sample multiplexing has been proposed as an experimental strategy to reduce sequencing-related batch effect (4–10), it does not control for technical biases introduced earlier during sample procuration and cell dissociation/sorting. In many situations, especially in clinical settings, it is difficult to “batch” the samples for library preparation. It is also often unclear, in single cell studies, what type of samples can serve as the best controls, nor how to make use of control samples during data integration. Thus, all current integration paradigms treat each sample as its own batch, and for studies that include “control” or “baseline” samples, the current standard is to ignore this information during the integration step and integrate cells across all samples in a way that is agnostic to experimental design.

There has been enormous progress on the problem of single cell data integration (11–21), highlighted by comprehensive reviews (21, 22). Despite this progress, key limitations remain in our current analysis paradigm, especially when faced with large-scale disease-focused single cell studies. The work in this paper is motivated by the following unresolved challenges: 1) Disease-focused studies are usually built on design principles such as case versus control cohorts and longitudinal sampling, yet neither current integration methods nor their benchmarks make use of these design principles to separate batch effects from biological variation. 2) We yet do not have a good grasp of how batch effects can vary across cell types and genes, and thus, current studies do not perform batch correction but rather, batch integration. Integration methods have tuning parameters that control the extent to which cells are aligned across samples to achieve uniformity, but without a clear understanding of how batch effects compare to biological variation, it is unclear how such parameters should be tuned. Different integration methods, and different parameter choices, often lead to different down-stream findings. Thus, current studies often take a black-box, trial-and-error approach to batch correction, compromising the reproducibility of their results. 3) Existing benchmarks have focused mostly on preservation of differences between cell types, but not the preservation of subtle cell type-specific changes between samples (21, 22). In many studies, the samples to be integrated are expected to differ biologically, and while the success of studies often hinge on the detection of subtle cell-type specific signals, current methods have been benchmarked mostly in the context of depletion/enrichment of major cell types.

In this study, we show that meaningful biological variation is unnecessarily removed in single cell data integration, and present a novel statistical framework, CellANOVA, that harnesses experimental design principles to explicitly quantify batch variation and recover the erased signals. Application of CellANOVA requires the choice of an existing integration method, as well as the identification of one or multiple control-pools: A control-pool is a set of samples whereby variation beyond what is preserved by the existing integration are not of interest to the study. The control-pool samples are utilized to estimate a latent linear space that captures cell- and gene-specific unwanted batch variations. By using only samples in the control pool in the estimation of the batch variation space, CellANOVA can recover any variation in the non-control samples that lie outside this space. Importantly, CellANOVA produces a batch corrected gene expression matrix which can be used for gene- and pathway-level downstream analyses. When applied with an existing integration, CellANOVA is fast and scalable to large single cell datasets.

## Results

### Defining Batch effects through construction of control-pool

We start by describing what is meant by “batch effect” in single cell studies, with a rigorous definition given in the next section. In single cell studies, each sample is a “batch of cells”, and we use the terms “batch” and “sample” interchangeably. It is unavoidable in high-plex experiments, regardless of protocol, for random technical variation to be introduced (23, 24). This technical variation, which can be specific to each cell and each gene, is confounded with biological variation in the observed data. Our definition of batch effects include, but are not limited to, such sample- and cell-specific technical variation.

The general use of the term “batch effect” has also included, sometimes explicitly (25) but often implicitly (26–28), biological variations that are deemed ignorable within the scope of a study. For example, consider a hypothetical study where one cannot control the time of day of sample collection. Circadian rhythms may affect our biological measurements, and if circadian effects are not of interest within the context of the study, then this biological variation can also be treated as a batch effect. In the statistical framework underlying CellANOVA, we make the vague concept of “batch effect” concrete through the construction of one or multiple control-pool(s), each consisting of a set of “control” or “baseline” samples that are not expected to differ from each other along the dimensions of interest to the study. We will include, as “batch effect”, any variation between the cells of these control-pool samples after conditioning on their latent cell state, which will be made precise in the next section. Thus, in addition to technical variation, batch effects can also include background biological variations within the control pool.

Given an existing best-effort integration of all samples, CellANOVA aims to recover any variation in the samples outside of the control-pool that may be erased during the integration. Variation among the samples in the control pool is used to learn the batch effect, and any variation that is orthogonal to the batch effect, as defined rigorously in the next section, can be recovered by CellANOVA.

Within this framework, the construction of the control sample pool is critical. To demonstrate this construction, we describe 4 studies of varying designs (Figure 1). The data from these studies will be used for illustration and benchmarking.

**Fig. 1:**
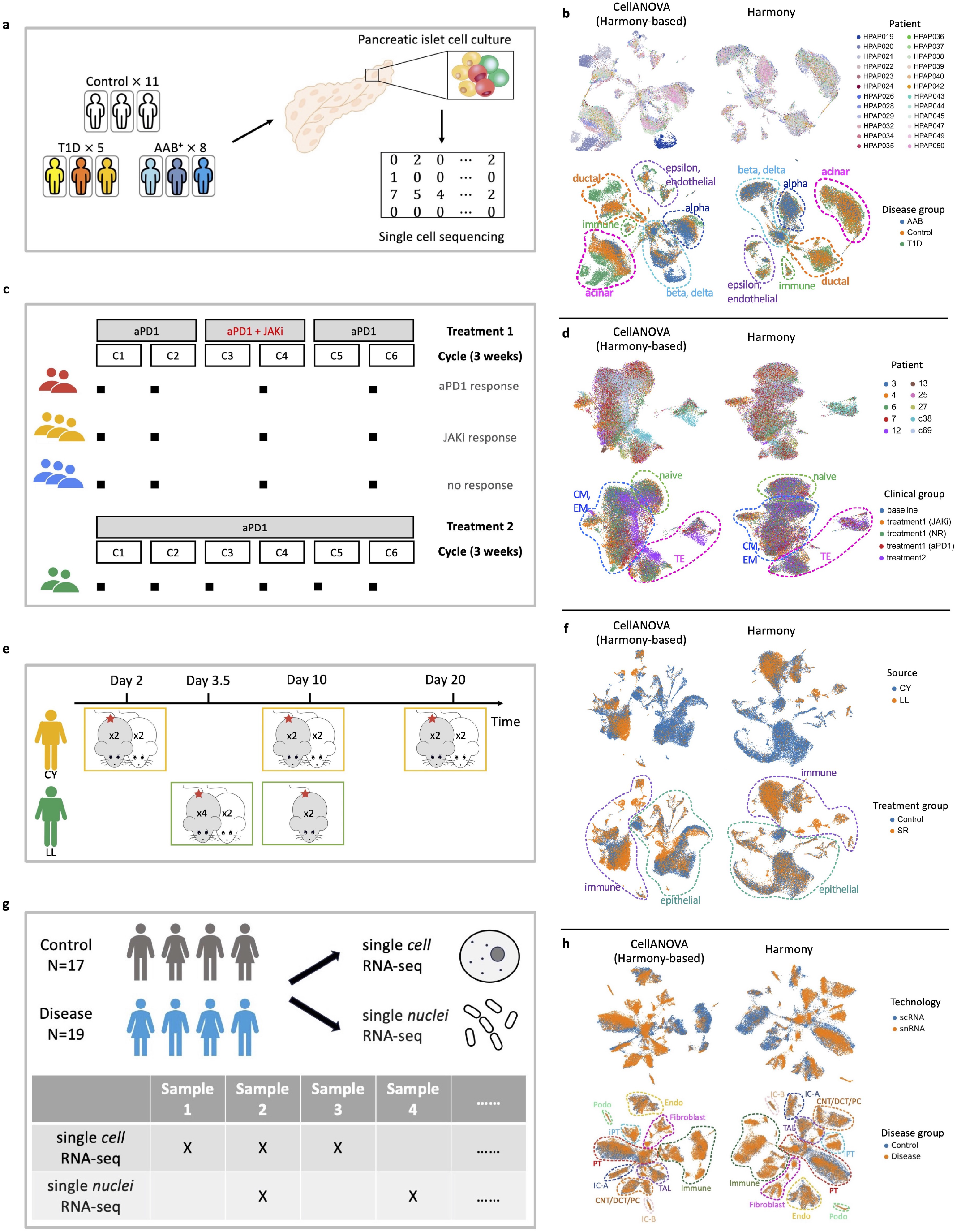
Examples of control-pool construction and integration results. (a) The case-control design in the type 1 diabetes (T1D) study involved 11 healthy individuals, 5 individuals with T1D, and 8 individuals with AAB+. The 11 healthy individuals are designated as the control pool.(c) The longitudinal design in the immunotherapy trial dataset involved 10 lung cancer patients undergoing 2 types of immunotherapy treatments sequenced at 4 time points. The 10 samples taken before treatment are designated as the control pool. (e) The irregular block design in the mouse radiation experiment performed by 2 technician teams, with a strong technician effect confounded with time. To separate time and treatment effects, we designate the 5 control samples as the control pool. (g) Case-control design of multimodal single-cell and single-nuclei RNA sequencing for kidney atlas-building study, with large batch effects from different technology platforms. We designate the 17 control samples (including samples from both scRNA-seq and snRNA-seq) as the control pool. UMAP visualizations of Harmony integration, with and without CellANOVA signal recovery, for each dataset. (b: type 1 diabetes study; d: immunotherapy trial dataset; f: mouse radiation experiment dataset; h: multimodal kidney dataset).

Example 1: Case-control design (Figure 1a). In this study of type 1 diabetes (T1D) from (29), cultured pancreatic islet cells from 11 healthy individuals, 5 individuals with T1D, and 8 individuals with no clinical presentation of T1D but positive for beta-cell auto-antibodies (AAB+) were sequenced. This is a common study design in clinical studies, where the goal is to identify disease-associated enrichment/depletion of cell types/states and cell-type specific differentially expressed genes. Clear batch effects are visible in the UMAP embedding of this data (Supplementary Figure 1), potentially confounding with disease status. Since the primary goal is to make comparisons between diseased and healthy individuals and between disease subgroups (i.e., T1D versus AAB+), we designate the 11 healthy individuals as the control-pool.

Example 2: Longitudinal design (Figure 1c). This is a study of 10 non-small cell lung cancer (NSCLC) patients undergoing two types of immunotherapy treatments, taken from (30). CD8 T cells sorted from peripheral blood were sequenced for each patient at four time points, including a baseline sample at time “0” right before the start of treatment. The patients differ by clinical outcome as well as by treatment regime. As described in the original study (30), eight of the patients received two cycles of pembrolizumab (aPD1), then itacitinib (JAKi for JAK inhibitor) concurrently with pembrolizumab for two cycles (Treatment 1). Then, pembrolizumab was continued until disease progression. Imaging was performed after the first two cycles of pembrolizumab and then after itacitinib (at the end of cycle 4) to assess tumor response. As shown in Figure 1c, patients in Treatment 1 were categorized into three groups: those who exhibited an early radiographic response to pembrolizumab by the end of cycle 2, whom we label aPD1 for “anti-PD1 blockade responsive”; those whose tumors failed to respond by the end of cycle 2 but responded at the end of cycle 4 with the addition of itacitinib, whom we label JAKi for “JAK-inhibitor responsive”, and those whose tumors remained refractory for the duration of treatment, whom we label NR for “non-responders”. With the scRNA-seq data, our goal is to examine how the CD8 T cells changed in these patients over the course of treatment, and how the T cell responses in these patients differ from those who were only treated with pembrolizumab (Treatment 2). (30). Although batch-effects seem less severe for this data (Supplementary Figure 2), as opposed to Example 1, the study aims to detect subtle changes in CD8 T cell expression that may require more finesse in integration. Since we are not interested in variation between the samples taken at baseline prior to treatment, we designate these 10 samples, one from each individual, as the control-pool.

Example 3: Irregular block design (Figure 1e). In this study on the effects of radiation on intestinal cells of mice, C56BL/6J mice were divided into a control group and a group that receives conventional radiation therapy. At days 2, 3.5, 10, and 20 post-irradiation, intestinal segments of two or more mice from each group were harvested and single cells were isolated and sequenced from the epithelial and lamina propria layers of the organ. The data analysis is complicated by the fact that two technician teams (whom we label CY and LL) performed experiments, and samples from the two teams are completely separated in the joint UMAP embedding (Supplementary Figure 3). The technician effect in this case is confounded with day: LL performed experiments on days 3.5 and 10, while CY performed experiments on day 2, 10 and 20. Control mice were included for most of the days, but not for LL on day 10. Since our goal is to quantify the time and treatment (radiation) effects on the cells, we designate the 5 control samples (2 from LL day 3.5, and 1 each from CY day 2, 10, and 20) as the control-pool.

Example 4: Case-control design with both single-cell and single-nuclei RNA sequencing (Figure 1g). In this study, Abedini et al. (31) collected kidney samples from *N* = 36 subjects, which were divided into two groups: (i) healthy controls (*N* = 17) and (ii) chronic kidney disease (*N* = 19), based on the estimated glomerular filtration rate and fibrosis. These samples underwent single-cell RNA sequencing (scRNA-seq) and/or single-nuclei RNA sequencing (snRNA-seq). The goal of this study is to construct an integrated atlas of healthy and fibrotic kidney to capture the changes associated with chronic kidney disease. However, the use of both single cell and single nuclei sequencing lead to significant batch effects (shown in Supplementary Figure 4), and biological variations between the patients may also confound the biological variations of interest. Since the objective is to mitigate batch effects resulting from different protocols while preserving the differences between the control and disease samples, we chose the 17 control samples, which includes both scRNA-seq and snRNA-seq samples, as our control pool.

In each of the above studies, the construction of the control-pool is a step of critical importance, as the control-pool samples serve not only as a biological baseline for comparison, but also as a representative sampling of the sources of unwanted variation which we will estimate using CellANOVA. Figure 1(h-j) previews the effects of CellANOVA when applied on a Harmony integration of each of the three datasets. Results when applied with other integration methods are shown in Supplementary Figures 1, 2, and 3. Visual examination reveals that the degree of inter-sample mixing varies substantially between integration methods, and in general, CellANOVA recovers a low-dimensional embedding that has more separation between disease states and treatment groups. Did CellANOVA effectively recover true and meaningful biological signal while keeping batches appropriately mixed? We start with an overview of the CellANOVA model and algorithm, followed by a detailed examination of its signal recovery capacity on these datasets.

### CellANOVA model and estimation procedure

Multi-sample single cell data comes in the form of *m* cell-by-feature expression matrices, **X**^(1)^,…, **X**^(*m*)^, where 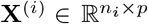 records measurements on *p* features (e.g., RNA expression levels) for *n*_*i*_ cells in the *i*th sample. We assume, without loss of generality, that samples *i* = 1, …, *m*_0_ are designated as the control-pool (Figure 2a). The case of multiple control pools is given in Methods. After modality-appropriate pre-processing (see Methods), we assume the data follows the following <u>cell</u> state space <u>a</u>nalysis <u>o</u>f <u>va</u>riance (CellANOVA) model:

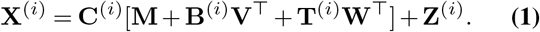

**Fig. 2:**
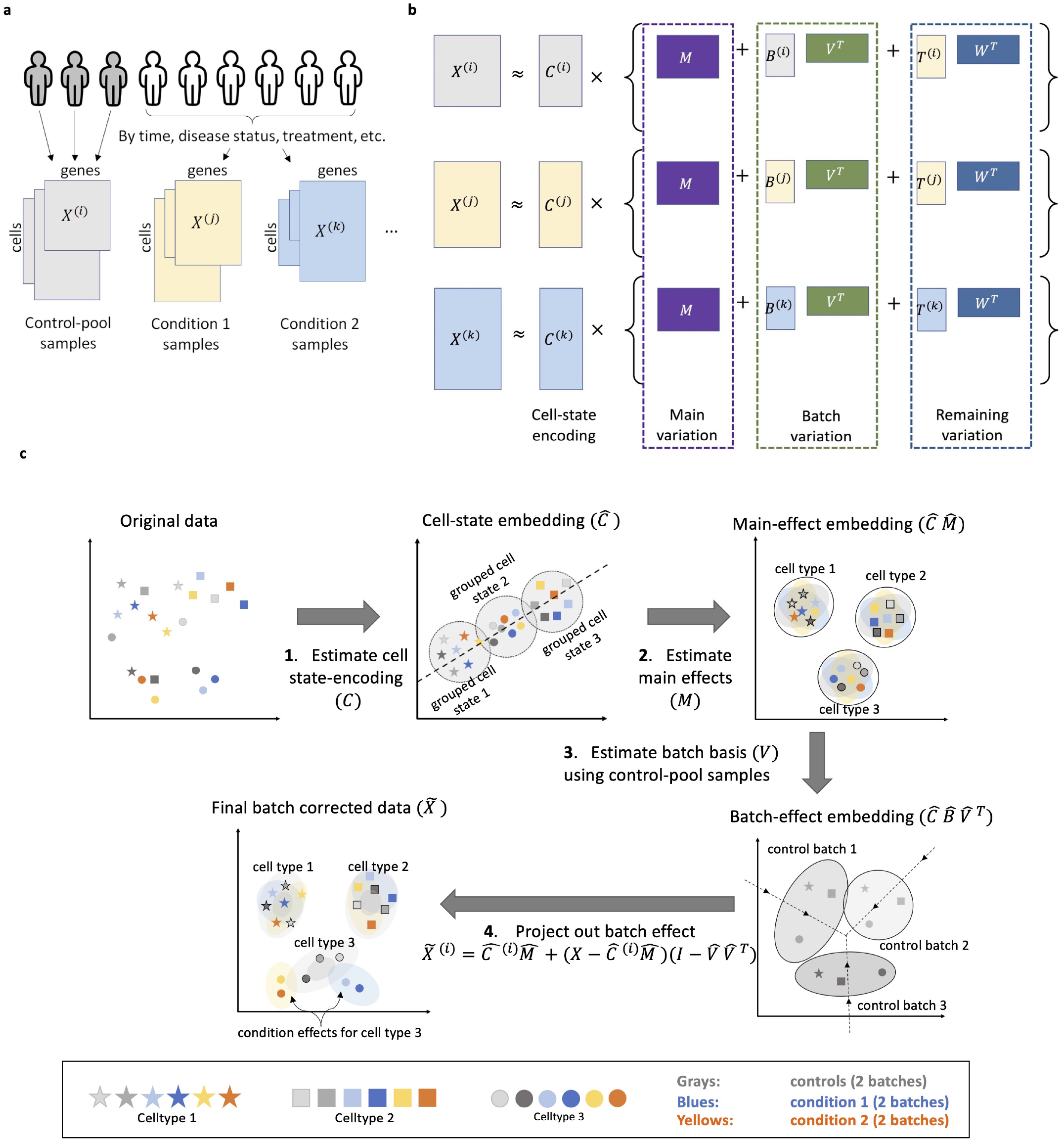
(a) “Pool-of-controls” design of multi-sample single-cell data. (b) The CellANOVA Model. (c) The CellANOVA algorithm. Step 1: Estimate cell state-encoding via singular value decomposition of an existing integration across samples. Step 2: Estimate main effects by regressing the original expression vectors on the cell state-encoding. Step 3: Estimate batch basis (**V**) using control-pool samples by performing singular value decomposition of the effect space after removing main effects. Step 4: Remove batch effects for all samples by projection into null space of **V**.

The model is shown in Figure 2b. Hereafter, for any matrix **A, A**^⊤^ stands for its transpose. In Eq. (1), 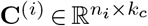encodes the unobserved state of each cell as a *k*_*C*_-dimensional vector, which we call the cell’s “state-encoding”. The cell state-encodings are multipled by a sum of three matrices: The main effect **M**, which captures average expression patterns across all samples in the dataset, the sample-specific batch effect **B**^(*i*)^**V**^⊤^, which captures unwanted variations that we wish to remove, and the sample specific signal matrix **T**^(*i*)^**W**^⊤^ which captures biologically meaningful variations that we wish to recover. Note that both the batch and signal matrices are products of sample-specific score matrices 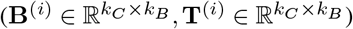 and cross-sample shared loading matrices 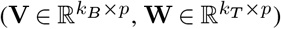. The matrix **V** can be interpreted as a basis of the linear space that captures the state-encoding-specific batch variations across samples. Since **V** is a key quantity in the CellANOVA algorithm, we give it the name “batch-basis matrix”. In contrast, **W** can be interpreted as a basis of the linear space that captures the remaining variation between samples after removing the batch variation. The last term, **Z**^*i*^, represents idiosyncratic noise that remains in the decomposition.

The identifiability constraints of the model and the details of the estimation procedure are given in Methods. To appreciate how CellANOVA builds on existing integration methods, we give an intuitive summary in Figure 2b. CellANOVA starts by applying an existing integration method to the entire dataset, to obtain an initial integrated data matrix across all samples. A singular value decomposition is performed on this integrated data matrix, and the cell state-encoding matrix (**C**) is estimated by the top *k*_*C*_ left singular vectors, which we denote by 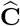 (step 1 in Figure 3c). Note that some biological differences between samples, such as enrichment/depletion of major cell types, are already preserved in **C**, as shown by extensive benchmarks of existing integration methods (21, 22). An embedding of **C** is what we commonly see in current integrated data embeddings.

**Fig. 3:**
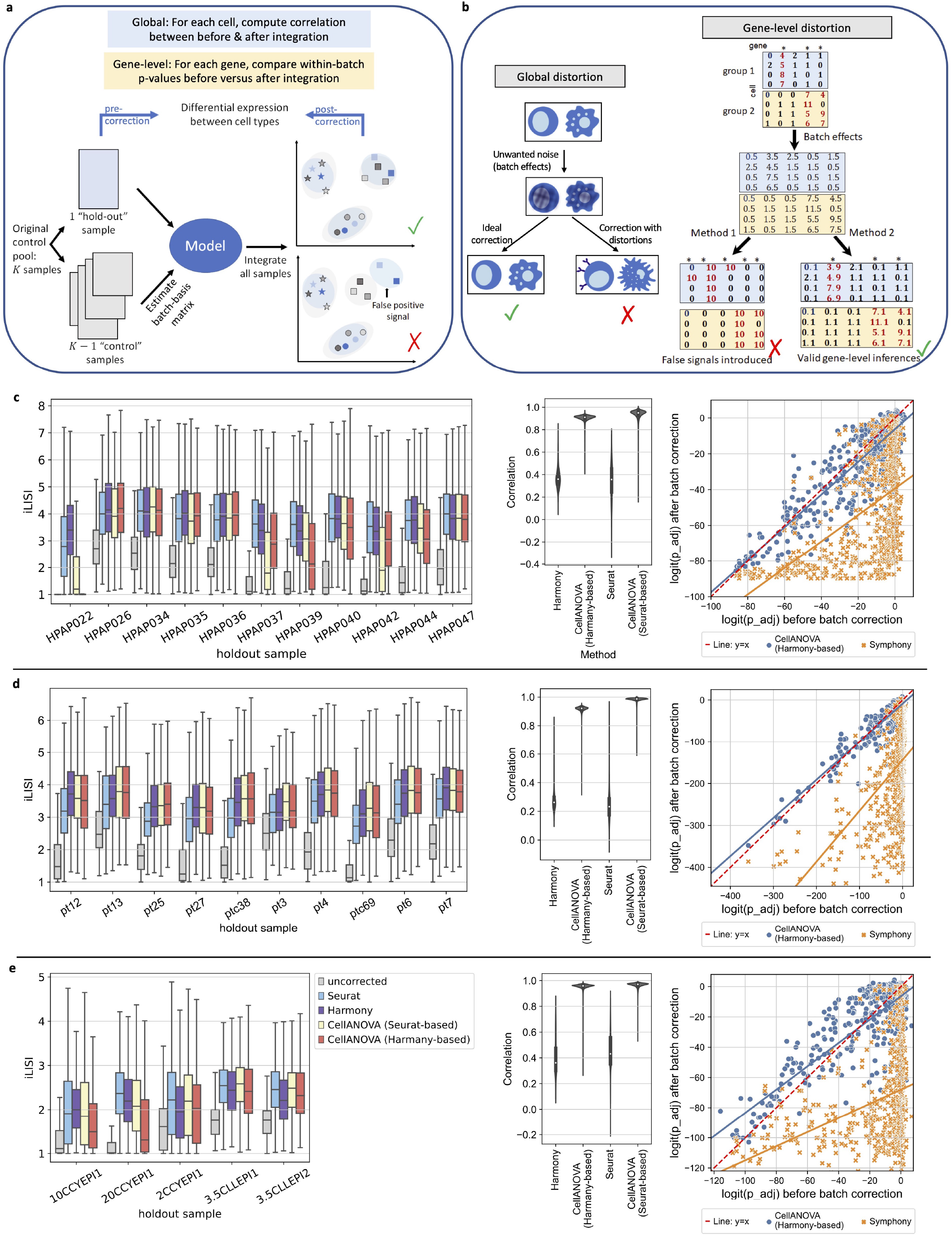
(a) Experiment workflow for benchmarking CellANOVA against existing state-of-the-art methods in removing unwanted batch variation, introducing global distortion (cell level) and gene-specific distortion (gene level). In each experiment run, we designated one control sample as a “fake” treatment sample (hold-out set) and used the remaining control samples to estimate the batch variation basis. On the hold-out sample, we performed DEG analysis using either uncorrected expression, or batch-corrected expression, between pre-defined cell types, obtaining a multiple-testing adjusted p-value for each gene for each comparison. We compute the correlation between pre- and post-expression for each cell. (b) Illustration of global distortion (left) and gene-specific distortion (right). Global distortion refers to the degree to which the integrated data differs from the original data prior to integration. Gene-specific distortion refers to the preservation of gene-level differences (or the lack thereof) between predefined cell groups. (c-e) Benchmark on type 1 diabetes dataset (c), immunotherapy trial dataset (d) and mouse radiation experiment dataset(e). LISI scores of the fake treatment sample after batch correction in each hold-out experiment are shown on the left. Correlations between pre- and post-CellANOVA correction gene expressions per cell are shown in the middle. Comparisons of p-values obtained from DEG analysis with or without CellANOVA correction are on the right.

Next, CellANOVA regresses the original data matrix **X**^(*i*)^ for each sample *i* on the estimated state-encoding of its cells, 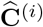, to obtain a matrix **R**^(*i*)^ of dimension *k*_*C*_ *× p*. An estimate of the main variation **M** is derived by averaging **R**^(*i*)^ across all samples (step 2 in Figure 2c). The batch-basis matrix **V** is estimated through quantifying the variation of **R**^(*i*)^ in only the control samples (i.e. centering **R**^(*i*)^ within the control-pool, followed by an SVD of a row-stacking of the centered matrices). In this way, the estimated batch-basis matrix 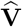 captures cell state-encoding specific variation between the control samples, post integration (step 3 of Figure 2c).

Although the batch-basis matrix is shared across all samples, CellANOVA gives an explicit estimate of the batch effect for each gene in each cell, in the form of 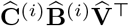. Thus, the batch effect for each cell is allowed to depend on its state-encoding (through **C**^(*i*)^) as well as its sample-of-origin (through **B**^(*i*)^). With data-derived estimates 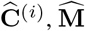, and 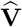 as described above, we remove the batch effect from sample *i* simply by projecting each of its cells, after encoding-specific centering, into the null space of 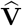 (step 4, Figure 2c),

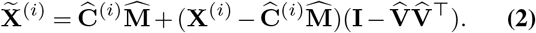

This projection gives a batch-corrected cell-by-feature matrix in the original data dimension, which can be used for downstream analysis such as differential expression, gene set enrichment, and trajectory reconstruction. For analyses such as clustering, it is necessary to start with a low-dimensional embedding. One can apply standard dimension reduction procedures to 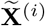. Note that such an embedding would differ from the embedding 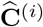 given by the initial integration, because of the additional variation 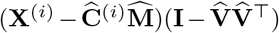 that has been recovered.

The CellANOVA model and estimation procedure also allow explicit delineation of the types of biological variation that can be recovered: to be separable from batch effects, the variation needs to lie outside the linear span of the batch-basis **V**, and only the component of the variation that is orthogonal to **V** can be recovered. CellANOVA is conservative in the sense that it removes any variation that is contained in the batch variation space estimated using the control-pool samples, which is, by design, to avoid introducing false positives.

There may be scenarios where there are multiple sets of control samples, where the variation between the samples within each control-pool are ignorable, but variation between samples belonging to different control-pools are of interest and should be retained. The CellANOVA model and estimation procedure can be adapted to this case, with details given in Methods.

### Is the integrated data free from batch effects?

CellANOVA uses only samples in the control-pool to estimate the batch-basis matrix **V**, which is then used to recover the biological variation between samples that might have been erased by existing integration methods. Hence, our first question is whether the variation recovered by CellANOVA is free of batch-effects. In other words, is the CellANOVA output as free of batch-effects, as compared to the original integration, especially for samples outside the control-pool that were not used in learning the batch-basis matrix? To answer this question, we devised the validation strategy shown in Figure 3a: For each dataset, we hold out one of the samples in the original control-pool and apply CellANOVA with the control-pool limited to the remaining samples. Then, we examine whether the hold-out control sample is effectively integrated with the other control-pool samples. Ideally, the hold-out control sample should be well-integrated with the other control-pool samples, even if it were not used in the estimation of the batch-basis matrix. This hold-out analysis can be viewed as a robustness test: For CellANOVA to be effective in removing batch effects from samples outside the control pool, the control-pool needs to exhibit the diversity of batch variations that affect all samples. If the holdout sample were not well-mixed with the other control-pool samples post-integration, then we would doubt that the batch-basis matrix estimated from the control-pool captures all of the unwanted variation in the study.

We applied this benchmarking strategy on the type 1 diabetes, immunotherapy trial, and mouse radiation datasets shown in Figure 1, The results for the kidney disease study, which involves the integration of single cell and single nuclei data, will be discussed separately in its own section. We compared CellANOVA (both Harmony- and Seurat-based) with Harmony (v0.1.1) (13), Seurat RPCA (v4.3.0) (32), LIGER (v1.0.0) (14, 33), Symphony (v0.1.1), and Seurat Reference Mapping (v4.3.0)(32). Harmony, Seurat, and LIGER were the three methods highlighted by (22). Symphony (34) is a reference mapping method that first constructs the reference atlas using Harmony and then maps queries into the same reference embedding. Seurat Reference Mapping (32) first constructs an integrated reference by applying the standard Seurat RPCA workflow. Then, it integrates the reference with the query data by correcting the query’s projected low-dimensional embeddings using the reference embedding as a template.

As proposed by (13), we used local inverse Simpson’s index (LISI) with respect to sample labels to measure the mixing of the hold-out sample with the remaining samples. The LISI of a cell is defined as the effective number of batches, properly scaled, within its *k*-nearest neighbors. We used *k* = 30. A higher value of LISI indicates more uniform batch mixing. Figures 3c-e (left panel) and Supplementary Figure 8 show the distribution of LISI scores across cells of the hold-out sample after integration, with the hold-out sample iteratively set to each of the 5 control-pool samples in the mouse radiation study, the 11 control-pool samples in the type 1 diabetes study, and the 10 control-pool samples in the immunotherapy trial study. Corresponding UMAPs showing alignment of the hold-out sample with the remaining control-pool samples is shown in Supplementary Figures 5, 6, and 7. Despite the fact that CellANOVA preserves more inter-sample variation (Figure 1h-j), the degree of mixing between the hold-out control sample and the rest of the control-pool is comparable to the original integration prior to signal recovery. This shows that, in recovering more signal, CellANOVA does not re-introduce unwanted variation.

### Does CellANOVA correct the distortion caused by integration?

The integration of cells across samples could inadvertently distort the data. Our goal is to remove batch effects while introducing minimal distortions, which is crucial to biological signal preservation and statistically sound downstream analyses. We consider two types of distortions: global and gene-specific (Figure 3b).

“Global distortion” refers to the degree to which the integrated data differs from the original data prior to integra-tion. While we certainly expect the integrated and original data matrices to differ, our goal should be to remove all of the unwanted variation while maintaining maximal possible similarity to the original data. Excessive global distortion reflects possible integration artifacts that could mislead downstream analyses. To assess the severity of global distortion, we computed Pearson’s correlation coefficient between each cell’s gene expression vectors before and after integration for each of the three datasets. A higher correlation indicates milder global distortion. We compared CellANOVA (both Harmony- and Seurat-based) to Harmony (v0.1.1), Seurat RPCA (v4.3.0), and Liger (v1.0.0). Although Harmony only outputs a low-dimensional embedding, we were able to extract a cell-by-feature matrix from the algorithm as described in Methods. Violin plots in Figure 3c-e (middle panel) and Supplementary Figures 9 (right panel) show that the cell-wise correlations between pre- and post-integration gene expression vectors were significantly improved by CellANOVA on all three datasets, *while* maintaining the mixing between batches (Figure 3c,d,e). Prior to signal recovery, the correlations between integrated and original data consistently averaged below 0.5. After signal recovery, the correlations increased to consistently average above 0.9. Interpreted in the context of the results in the previous section (Figure 3), we can conclude that CellANOVA substantially reduce global distortion, preserving the highest similarity to the original data while effectively removing technical and biological unwanted variation.

In addition to global distortion, we also consider “gene-specific distortion”, which refers to the preservation of gene-level differences (or the lack thereof) between predefined cell groups. The concern here is that integration may artificially strengthen or weaken differential expression signals between cell groups. Referring to Figure 3b, we expect differential expression signals in the observed data to be corrupted by batch effects. After integration, we would like to recover the true differentially expressed genes between cell populations, and avoid artificially inflating the significance of genes that are not truly differentially expressed.

To assess gene-level distortions, we developed the novel strategy shown in Figure 3a. First, we hold out one sample and use the remaining samples to fit the **M** and **V** matrices in CellANOVA. On the hold-out sample, we identify differentially expressed genes between pre-defined cell types, obtaining a multiple-testing adjusted p-value for each gene for each comparison. We then compute batch-corrected expression values for the hold-out sample using the fitted model and perform post-correction differential expression analysis between cell types, again obtaining adjusted p-values for each gene. Note that the pre- and post-correction p-values are computed using exactly the same sets of cells, all derived from the same sample (batch). Since they all come from the same batch, the pre-correction differences between these cells are not confounded by batch effects, and thus, the post-correction p-values should resemble their pre-correction counterparts. Figure 3c-e (right panel) plot the pre- and post-correction adjusted p-values against each other, with a high correlation indicating lower gene-wise distortions. For a fair comparison, we compared our approach against Harmony integration with Symphony mapping, which, like CellANOVA, treats the hold-out sample as a query and the remaining samples as the reference. In this way, the hold-out sample is not used in fitting the model. The scatter plots in Figure 3c-e show that p-values obtained in the differential expression analysis using CellANOVA-integrated data are highly correlated with those obtained prior to integration, indicating minimal gene-level distortions. Importantly, this analysis shows that CellANOVA maintains valid p-values post integration. In contrast, current integration methods artificially reduce the p-values, making it difficult to control type 1 error in downstream comparisons between cell types.

### Does CellANOVA recover true and meaningful biological differences between samples?

CellANOVA is motivated by the desire to recover biologically meaningful variation which may be erased during single cell data integration. We now explicitly evaluate the extent of biological signal recovery, focusing specifically on the recovery of meaningful differences between samples. Despite the growing knowledge of *intra*-sample cellular variation (e.g., what are the cell types and the differences between them), studies usually have little a priori knowledge of what comprises true *between*-sample variation beyond possibly a small set of positive controls. Thus, we employ two strategies for assessing the extent of recovery of true between-sample differences: (1) The leveraging of meaningful sample groupings, and (2) the leveraging of cross-modality and cross-study replication. We first focus on strategy (1).

In most single cell studies, samples are labeled in meaningful ways such as by disease status, treatment arm, or collection time. We use “condition” as a generic term to refer to a grouping of samples based on a predefined label. Comparisons between conditions are usually of primary interest to a study, as it is for the four studies in Figure 1: In the type-1 diabetes and kidney disease study, the goal is to compare between disease states; in the immunotherapy trial, the goal is to compare between different treatment-response groups, and within each group, across time; In the mouse irradiation study, the goal is to compare between the treatment and control arms, and within the treatment arm, across time. Figure 4a shows a toy example of two conditions, each comprising two samples. An overly aggressive integration might completely intermix all samples (left panel), erasing not only batch effects but also meaningful differences between conditions. On the other hand, an integration could fail to completely remove batch effects (right panel), the presence of which would confuscate downstream comparisons between conditions. An effective integration should remove batch differences while preserving those differences between conditions that lie outside of the span of the batch-basis **V** (middle panel).

**Fig. 4:**
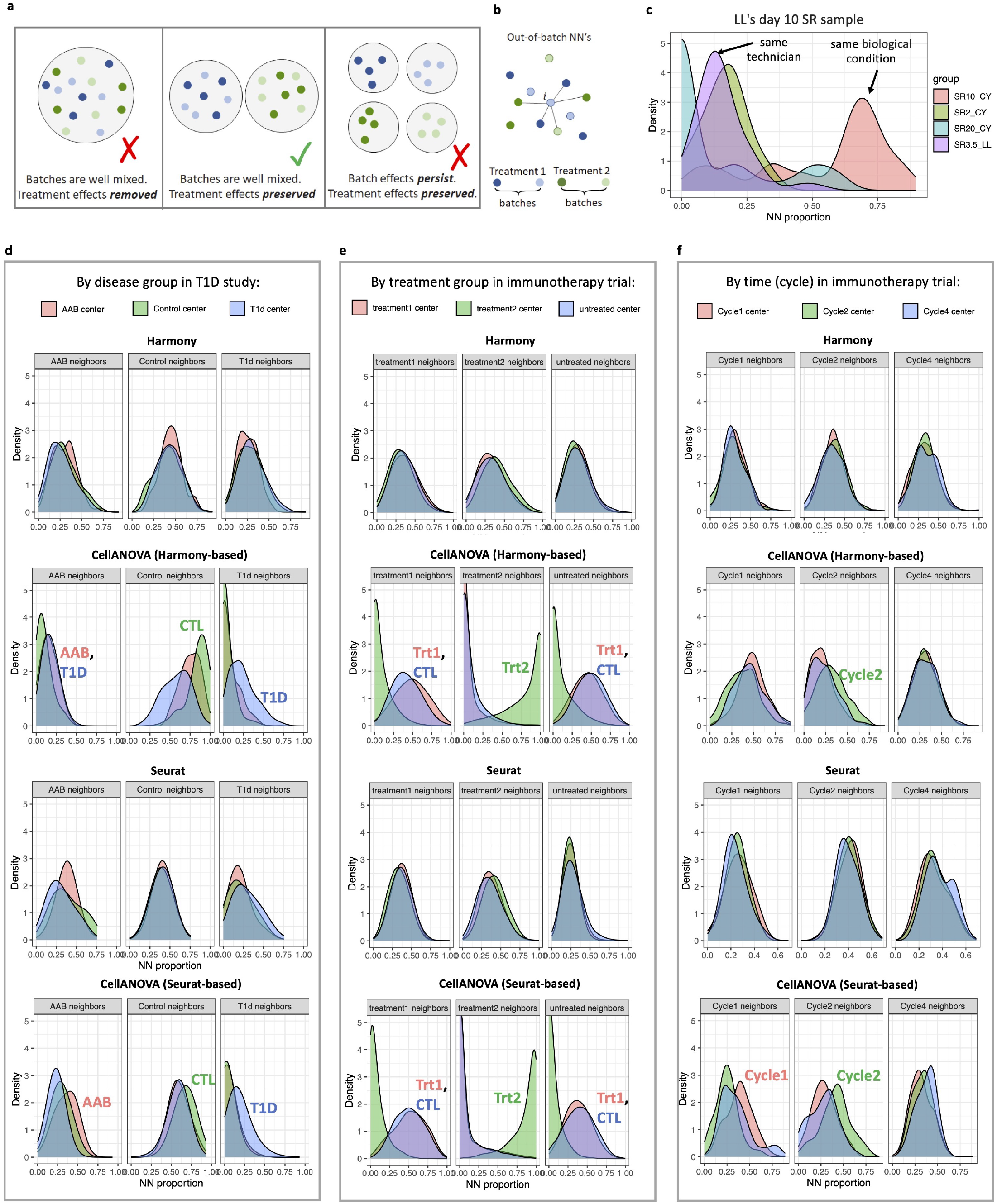
(a) Illustration of batch integration with signal preservation. An effective integration removes batch differences while preserving differences between conditions (middle). An overly aggressive integration erases meaningful differences between conditions (left). An ineffective integration fails to remove batch effects (right). (b) Illustration of out-of-batch nearest neighbors. We search for a cell’s nearest neighbors, only cells outside of the cell’s own batch (i.e., sample) are considered. NN: nearest neighbor. (c) Nearest-neighbor composition for cells from Leo’s day 10 SR sample, after integration by CellANOVA. The enrichment of cells from the same biological condition rather than the same technician indicates effective batch removal with signal preservation. (d-f) Benchmarking signal preservation after batch correction on three datasets using out-of-batch nearest neighbor proportion: (d) ductal cells in the T1D dataset, (e) all CD8 T cells from all patient groups in the immunotherapy trial dataset, (f) non-naive CD8 T cells from the JAKi group in the immunotherapy trial dataset. Enrichment of cells from the same treatment condition indicates the recovery of biological differences specific to the condition and cell type.

To examine the extent of post-integration recovery of meaningful variation between samples, we devised the following strategy: First, for each cell, its thirty out-of-batch nearest neighbors are identified within the integrated cell embedding (Figure 4b). “Out-of-batch” means that when we search for a cell’s nearest neighbors, only cells outside of the cell’s own batch (i.e., sample) are considered. If the cell belongs to a cell state where there are differences between conditions, we expect these out-of-batch neighbors to be enriched for cells from the same condition. Thus, for each cell we can compute the proportion of its out-of-batch nearest neighbors that come from each condition. Ideally, this density should be highest for the condition that matches the condition of the center cell. Since the condition label is not used by CellANOVA in signal recovery, any same-condition enrichment (i.e. density shifting to the right) is proof that biological signals that separate the given condition from the rest has been recovered in the integration.

As proof-of-principle, consider the mouse irradiation dataset (Figure 1e), which was generated by two different technicians, CY and LL. As shown in Supplementary Figure 3f, before integration, we observed that technician difference was the main source of batch effects. CY sequenced samples from day 2, day 10, and day 20, whereas LL sequenced samples from day 3.5 and day 10. At each time point, samples sequenced by CY included a set of control mice (denoted by C) and a set of irradiation-treated mice (denoted by SR), while LL included two sets of controls and four sets of SR samples at day 3.5, and no controls at day 10. Applying CellANOVA, we used the 5 control samples to estimate the batch-basis matrix and then removed batch effects for all samples. Figure 4c shows the nearest-neighbor composition for cells from LL’s day 10 SR sample. After CellANOVA integration, cells from LL’s day 10 SR sample are mostly surrounded by cells from CY’s day 10 SR sample, the sample to which it should be biologically most similar. This enrichment of cells from the same biological condition (i.e. the same number of days after treatment) rather than the same technician indicates preservation of biological signals after integration. Importantly, CellANOVA successfully matched LL’s day 10 SR sample to its correct counterpart among CY’s samples, even though there were no control samples from day 10 for LL. This indicates that, for estimation of the batch latent space, it is not necessary to include controls within every biological condition as long as the control-pool samples are sufficiently representative for capturing potential batch effects.

Now consider the type 1 diabetes and immunotherapy trial datasets, with the kidney disease dataset discussed in its own section later. In the type 1 diabetes (T1D) study (29), it was observed that a family of ductal cells separated into T1D-enriched and control/AAB-enriched subpopulations. Analyses conducted by (29) suggested that this separation was driven by true transcriptomic differences rather than technical biases. To examine whether CellANOVA corroborates this finding, Figure 4d plots the neighborhood composition distributions of ductal cells from the Control, AAB, and T1D conditions, after integration by Harmony- or Seurat-based CellANOVA and initial integration by Harmony and Seurat, respectively. We observe that, after CellANOVA signal recovery, strong differences in the ductal cells are observed between the T1D, AAB, and control conditions, as the out-of-batch nearest neighbors of ductal cells are enriched for cells of the same condition, regardless of whether CellANOVA was applied to Harmony- or Seurat-based integrations. Furthermore, AAB neighbors were enriched in both AAB and T1D groups, while control cells were enriched in control and AAB groups. These findings suggest that AAB is an intermediate state between T1D samples and healthy samples, which aligns with our current understanding of type 1 diabetes. In contrast, without CellANOVA signal recovery, the differences in ductal cells between the three groups was mostly erased by integration as shown in Figure 4d.

The immunotherapy trial dataset can be stratified by treatmemet: before treatment (baseline at cycle 1), after treatment 1 (pembrolizumab + itacitinib), and after treatment 2 (pembrolizumab only). Additionally, the samples collected after either treatment 1 or treatment 2 were sequenced at multiple time points, and thus can be stratified by time after treatment. In Figure 4e, we first grouped cells based on treatment and plotted the out-of-batch nearest neighbor composition for all CD8 T cells. After application of CellANOVA (either Harmony- or Seurat-based), the samples collected post treatment 2 are separated from baseline and from the samples collected post treatment 1. In contrast, the differences between treatments were not visible from the initial integration by Seurat or Harmony. We see similar trends when stratified by time: Figure 4f shows the out-of-batch nearest neighbor composition for non-naive CD8 T cells from the JAKi-responsive subjects, as defined in the original study. Mathews et al. (30) reported that one defining characteristic of JAKi responsive patients was that their tumors were still growing in cycle 2 but shrinks at cycle 4. Our analysis shows that, for these patients, CD8 T cells from cycle 2 are distinct after CellANOVA recovery, while this difference is absent in the initial integration. This is true whether the initial integration is by Seurat or Harmony. This cycle effect in the immunotherapy trial will be further compared to sample-matched flow cytometry in the next section.

### CellANOVA recovers subtle signals confirmed by matched flow cytometry

To further evaluate the extent of biological signal recovery and the effect of distortion correction on downstream analyses, we consider longitudinal changes in the NSCLC immunotherapy trial scRNA-seq data (Figure 1c), and compare our findings to those made by flow cytometry for the same samples. We first considered the longitudinal molecular signature reported by (30) for circulating CD8 T cells in this patient cohort: By measuring the proliferation marker Ki67 in these same patients at the same time points, Mathew et al. (30) showed that the anti-PD1 responders have a statistically significant increase in Ki67+ non-naive CD8 T cells between cycle 1 and 2, whereas the patients in the JAKi group, who did not respond by cycle 3, lacked this initial increase in non-naive CD8 T cell proliferation (shown in Figure 5a, lower panel). This is concordant with biological intuition, as PD1 blockade should reactivate anti-tumor CD8 T cells and thus stimulate proliferation, at least in the aPD1 responsive patients. To examine non-naive CD8 T cell proliferation in the scRNA-seq data, we applied CellANOVA, treating all samples from cycle 1 (before the start of pembrolizumab) as the control-pool in the estimation of the batch variation basis, to integrate samples from all of the cycles (1, 2, 4, and 6). After integration, we performed differential expression analysis followed by pathway-enrichment analysis between consecutive sampling times (see Methods). We focused on the G2-M Checkpoint pathway in the Molecular Signature Database (MSigDB) (35), a key pathway in cell division and a crucial component of the cell cycle. As shown in Figure 5a (upper panel), gene set enrichment analysis (GSEA) on CellANOVA-integrated data shows significant enrichment for the G2-M Checkpoint pathway (p-value *<* 0.05 using Harmony-based CellANOVA, p-value *<* 0.1 using Seurat-based CellANOVA) within the aPD1 group from cycle 1 to cycle 2. Within the aPD1 group, activity in this pathway dropped from cycle 2 to cycle 4, as indicated by a significant negative enrichment score (p-value *<* 0.01 using both Harmony- and Seurat-based CellANOVA). These results corroborate the transient proliferative burst in non-naive CD8 T cells in aPD1 patients identified by flow cytometry. For patients in the JAKi group, CellANOVA found no significant proliferation burst in non-naive T cells within the first 4 cycles, which is also consistent with flow cytometry results. In contrast, patterns in the G2-M Checkpoint pathway after the initial integration by Harmony, Liger and Seurat were not consistent with each other, nor with flow cytometry results, nor with biological intuition (Figure 5a, upper panel).

**Fig. 5:**
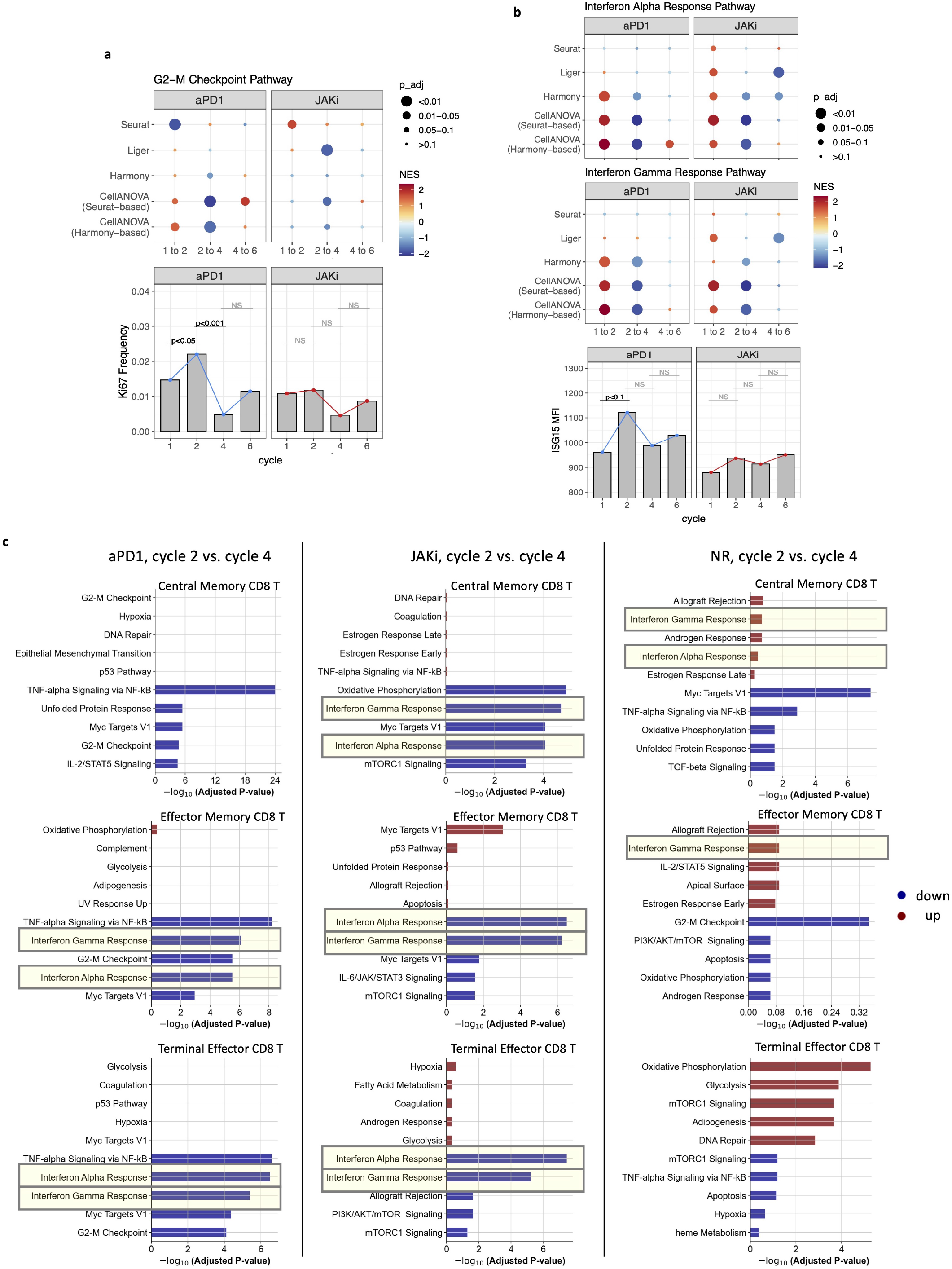
Comparison of pathway enrichment analysis based on scRNA-seq versus flow cytometry of corresponding markers in NSCLC immunotherapy trial data. (a) G2-M checkpoint pathway enrichment (scRNA-seq) versus Ki67 frequency (flow cytometry). A positive normalized enrichment score (NES) from GSEA indicates higher pathway enrichment in the later time points. Both Ki67 and G2-M checkpoint pathway activity measure cell proliferation. (b) Interferon alpha/gamma response pathway enrichment (scRNA-seq) versus ISG15 mean fluorescence intensity (flow cytometry). (c) Cell-subtype-specific gene set analysis within each response group between cycle 2 and cycle 4 after Harmony-based CellANOVA integration. Top 5 up-regulated and down-regulated pathways in cycle 4 compared to cycle 2 are shown.

Next, consider longitudinal changes of two key signalling pathways over the course of the treatment, namely the Interferon Alpha Response pathway and the Interferon Gamma Response pathway. As described in the original study (30), we anticipate a higher enrichment of both pathways in cycle 2, as compared to cycle 1, due to the chronic inflammation observed in cancer patients. Conversely, the addition of itacitinib, a JAK1 inhibitor which suppresses JAK1-dependent cytokine signaling like interferon, starting from cycle 3 should cause a decrease in the activities of both pathways in cycle 4 as compared to cycle 2. Mathews et al. (30) detected this longitudinal change using flow cytometry data: Figure 5b (lower panel), shows that the mean fluorescence intensity (MFI) for protein ISG15 (a direct readout from both interferon pathways) increased from cycle 1 to cycle 2, and then decreased from cycle 2 to cycle 4. Focusing on the aPD1 and JAKi group, as shown in Figure 5b (upper and middle panel) and Supplementary Figure 10, the GSEA results using CellANOVA-integrated data demonstrated the initial enrichment of both pathways in cycle 2, followed by their suppression in cycle 4. This is concordant with both the flow cytometry results and our knowledge of the effects that JAKi should have. In the initial design-blind integration by Harmony, Liger, and Seurat, longitudinal changes in both interferon response pathways are not consistent with the flow cytometry data.

Next, we used CellANOVA-integrated data to investigate the cell-type specific gene expression changes that may shed light on why the patients in the aPD1 and JAKi group showed tumor regression while those in the NR group did not. Focusing on cycle 2 and cycle 4, we performed gene set analysis on each subtype of non-naive CD8 T cells, including central memory, effector memory, and terminal effector cells, and identified the top 5 up-regulated and down-regulated pathways in cycle 4 for each cell subtype per patient group. Our analysis revealed that the activity of interferon-related pathways decreased from cycle 2 to cycle 4 in the aPD1 and JAKi groups, but not in the non-responders (Figure 5c, Supplementary Figure 11). This suggests that JAKi did not take effect in the non-responders, which may contribute to continued inflammation in these patients and worse outcomes. This drop in activity of interferon-related pathways is especially significant in the more differentiated cell subtypes (terminal effector and effector memory CD8 T), but is also significant in the central memory cell CD8 T cells for patients in the JAKi responsive group. For the non-responders, interferon-related pathways actually have a significant increase in activity according to the CellANOVA recovery based on Seurat integration (Supplementary Figure 11). These findings confirm the importance of temporal modulation of the interferon-stimulation response in immunotherapy.

### CellANOVA recovers replicable signals in single cell and single nuclei data integration

The integration of single-cell RNA sequencing (scRNA-seq) and single-nuclei RNA sequencing (snRNA-seq) data has been well appreciated to be challenging, as these two protocols are measuring two different RNA populations within each cell. To overcome the gross differences between these two protocols, data integration often incurs substantial distortion and signal loss. We consider data from a recent study (31) on the human kidney, with cells collected from 17 healthy individuals and 19 individuals diagnosed with chronic kidney disease (CKD). The goal of this study was to construct a comprehensive atlas of cell types in the kidney, both in its healthy state as well as during CKD. Samples from each individual were sequenced using scRNA-seq, snRNA-seq, or both protocols.

We applied CellANOVA based on an initial Harmony integration of the data. We chose to start with a Harmony-based integration because Harmony was able to integrate the data to achieve a better mixing of the single cell and single nuclei samples within the major cell types of the kidney, as compared to other methods. However, the extent to which Harmony’s integration erased disease-relevant signals is not clear, and the objective of CellANOVA is to recover any such lost signals. We first verify that CellANOVA does not re-introduce batch effects while correcting the distortions in this challenging data integration task. Figure 6a (left panel) shows the distribution of iLISI scores across cells of each hold-out sample after integration, with the holdout sample iterating across the samples in the control pool as according to the scheme in Figure 3a. We see that CellANOVA achieves higher iLISI scores compared to Harmony, indicating better batch mixing, despite the fact that it adds inter-sample variation back to the Harmony-integrated data. Figure 6a (right panel) shows the distortion metrics described in Figure 3 on this data: CellANOVA corrects both global (cell-wise) and gene-level distortion introduced by Harmony integration. Thus, we can conclude that CellANOVA improves batch mixing while correcting data distortions.

**Fig. 6:**
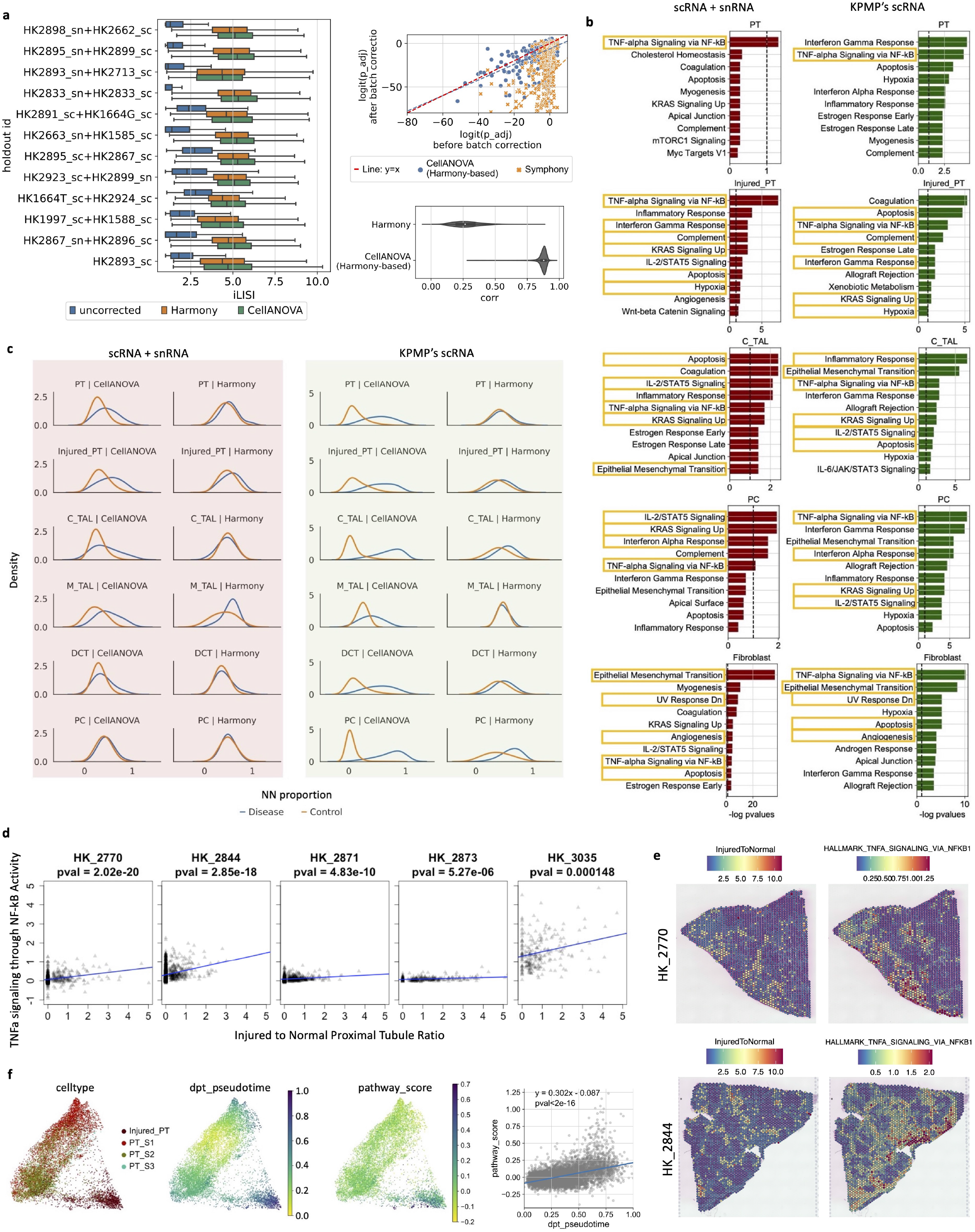
Evaluation of CellANOVA in motimodal data integration. (a) Assessment of CellANOVA in batch removal and distortion correction. Left panel: distribution of iLISI scores across cells of each hold-out sample based on unintegrated data, Harmony-integrated data, and CellANOVA-integrated data. Top-right panel: comparisons of p-values obtained from DEG analysis with or without CellANOVA. Bottom-right panel: correlations between pre- and post-CellANOVA correction gene expressions per cell. (b) Top ten upregulated pathways identified within the disease condition for each specific cell type in the Abedini et al. data and the KPMP data, using batch-corrected expression data generated by CellANOVA. (c) Density plot for the distribution of out-of-batch nearest neighbor proportion from disease or control conditions around diseased cells. (d) Scatter plots of TNF-alpha signaling via NF-kB pathway activity score versus injured proximal tubule to normal proximal tubule cell ratio for each Visium slice, with p-values calculated from linear regression. (e) Spatial distribution of spot-specific injured proximal tubule to normal proximal tubule cell ratio (left) and spatial distribution of the activity score of TNF-alpha signaling via NF-kB pathway (right) on Visium slice from sample HK_2770 (top) and HK_2844 (bottom), respectively. (f) Left panel: UMAP visualization of CellANOVA-integrated PT and iPT cells, with cells colored by cell type, diffusion pseudotime from PAGA trajectory analysis, and pathway activity score of TNF-alpha signaling via NF-kB. Right panel: scatter plot of diffusion pseudotime along the trajectory versus TNF-alpha signaling via NF-kB pathway activity score.

Next, we evaluate the effectiveness of CellANOVA in recovering biological differences between the disease and normal samples. For an unbiased, global assessment, we quantified the overlap of the identified differences in the Abedini et al. data with those identified in the Kidney Precision Medicine Project (KPMP) by Lake et al.(36), which is an independent study of the same tissue and disease. From the KPMP study, we took only the scRNA-seq data, which consists of 20 normal and 15 chronic kidney disease samples, and integrated them using Harmony followed by CellANOVA signal recovery. Since we only used scRNA-seq data from KPMP, we are confident that the signals thus identified are devoid of single cell to single nuclei integration artifacts, and thus signals identified in the Abedini et al. (2023) data and replicated in this study are more likely to be real. We focused on the kidney cell types with high enough cell count: Proximal tubule cells (PT), Injured proximal tubule cells (Injured-PT), cells of the cortical and medullary thick ascending loop of Henle (C-Tal and M-Tal), cells of the distal convoluted tubule (DCT), principal cells of collecting duct (PC), intercalated cells, podocytes, connecting tubule (CNT), and fibroblast. Within the disease condition, we computed the proportion of each cell’s out-of-batch nearest neighbors originating from either the disease or control samples, respectively, following the scheme of Figure 4a,b. As shown in Figure 6c, CellANOVA substantially increased the enrichment of diseased cells among the out-of-batch nearest neighbors of diseased cells for most cell types. This can be seen in the neighborhood density plots as well as the total variation statistic (Supplementary Figure 12) This enrichment is replicated in the independent KPMP scRNA-seq data. In contrast, without CellANOVA signal recovery, for many cell types there is no detectable separation between the normal and disease samples. These findings collectively suggest systemic changes in almost all of the major cell types in the kidney during CKD.

To elucidate the cell-type specific genes and pathways involved in CKD, we next performed differential gene expression analysis across the major kidney cell types, followed by pathway enrichment analysis. We used the batch-corrected expression data generated by CellANOVA to ensure that our analysis was devoid of any confounding batch effects. The results, before and after CellANOVA signal recovery, are presented in Figure 6b and Supplementary Figure 13. Figure 6b shows the top ten upregulated pathways identified within the disease condition for each specific cell type in the Abedini et al. data and the KPMP data. Here, again, we use the KPMP data to benchmark the replicability of the findings, with the replicated pathways highlighted in Figure 6b. CellANOVA enabled the detection of many cell-type specific disease-activated pathways, the majority of which are replicated in the KPMP data. In contrast, prior to CellANOVA signal recovery (Supplementary Figure 13), few significant disease-activated pathways were identified.

Now, consider two specific pathways that showed the strongest cell type-specific activation in the Abedini et al. data: TNF-alpha signaling via NF-kB in proximal tubule cells, and epithelial mesenchymal transition in fibroblasts. Both signals are replicated in the KPMP data. Prior to CellANOVA signal recovery, the TNF-alpha signaling via NFkB signal was not identified in neither the Abedini et al. nor the KPMP data, and the epithelial mesenchymal transition signal was only identified in the Abedini et al. data, and not in the KPMP data. To follow up on these two cell-type specific pathway activation results, we analyzed the spatial transcriptomic (VISIUM) data from 5 CKD kidney slices in Abedini et al. Since the generation of spatial transcriptomic data does not involve cell dissociation, the spatial transcriptomic data should be devoid of the dissociation-related batch effects that plague single cell sequencing data. Thus, spatial localization of pathway activity to spots where a given cell type is enriched provides independent evidence that the pathway is active in the given cell type in CKD tissue. First, consider TNF-alpha signaling via NF-kB, which our analysis suggests should be activated in proximal tubule cells during injury. Indeed, for each of the five slices, the activity of TNF-alpha signaling via NF-kB localizes to regions of the tissue with a high injured proximal tubule to normal proximal tubule cell ratio, with p-value *<* 10^−3^ in all five slices (Figure 6d). Figure 6e shows this colocalization pattern on two VISIUM slices with the most significant p values. To further investigate the enrichment of TNF-alpha signaling via NF-kB in regions exhibiting a high ratio of injured to normal proximal tubule cells, we employed the CellANOVA output to perform trajectory analysis and computed TNF-alpha signaling via NF-kB pathway activity score for both PT and iPT cells along this trajectory. As shown in Figure 6f, our analysis reveals a significant positive correlation (p-value *<* 10^−16^) between TNF-alpha signaling via NF-kB pathway activity and the transition from PT to iPT. As proximal tubule cells become injured, TNF-alpha signaling via NF-kB pathway is activated. Similarly, we expect the epithelial mesenchymal transition pathway to be activated in fibroblasts in disease tissue, and indeed, the activity of this pathway significantly colocalizes with fibroblasts in all of the five slices (Supplementary Figure 14).

### Benchmarks by simulation and cell type hold-out experiments

To better understand what types of signals can be recovered by CellANOVA and the extent of signal recovery, we used cell-type hold out experiments and simulations for a systematic comparison of methods.

We first examine whether a condition-specific cell type that is only subtly distinguished from other shared cell types can be preserved during integration. While existing studies have extensively benchmarked the preservation of cell type enrichment/depletion signals (see, e.g. (22)), they focused mostly on cell types that are well separated in low dimensional embeddings were considered. To test on subtly distinguished cell subtypes, we used the NSCLC immunotherapy trial dataset, for which the goal of the original study (30) was to identify condition-specific CD8 T cell subtypes. We will consider four CD8 T cell subtypes: naive, central memory, effector memory, and terminal effector cells. We took samples from baseline time points and artificially divided them into a treatment group (2 samples) and a control group (remaining 8 samples). Then we removed the terminal effector CD8 T cells from the control samples so that this became a treatment-specific cell type. We used samples in the control group to estimate the batch variation basis and then corrected batch effects for both control and treatment samples. As shown in Supplementary Figure 15a, CellANOVA successfully recovers the separation of the treatment-specific cell type (terminal effector CD8 T) in the UMAP space. On the contrary, the initial integration erases this subtle signal, mixing the terminal effector with effector memory CD8 T cells. We also used LISI scores to quantitatively evaluate the removal of batch effects versus the preservation of signal, shown in Supplementary Figure 15b. On the left, for the control samples, CellANOVA achieves comparable LISI scores as existing methods, indicating that CellANOVA effectively removes batch effects. On the right, the LISI scores for the treatment-specific terminal effector CD8 T cells are much lower after CellANOVA signal recovery, as compared to in the initial integration, indicating that CellANOVA recovered this cell type. Since the LISI score measures the effective number of batches in the neighborhood of each cell, and the terminal effector CD8 T cells are only present in two batches, the ideal LISI score of this treatment-specific cell type should be no larger than two.

Next, consider a scenario where the treatment does not introduce new cell types but alters the expression level of genes in existing cell types. As shown in Supplementary Figure 15c, we simulated a dataset with six cell types (CT1, CT2, …, CD6), across five control batches and two treatment batches. We introduced differentially expressed genes by increasing the expression of a set of genes in CT6 cells in treatment batches, see Methods for details of simulation model. In Supplementary Figure 15c, UMAP plots demonstrate that only CellANOVA recovers the within-cell-type (CT6-specific) differences between control and treatment groups, while this subtle difference is lost in the initial integration. The heatmaps of the batch-corrected expressions of differentially expressed genes between control and treatment groups are shown in Supplementary Figure 16, which confirms that CellANOVA recovers such subtle cell-type specific differential expression signals. To further evaluate the efficacy of CellANOVA, we used the batch-corrected expressions of CT6 cells to perform differential expression analysis between control and treatment groups, and the resulting ROC curves of different methods are shown in Supplementary Figure 15e, where CellANOVA significantly improves the integration results of Seurat and Harmony.

### What comprises batch effects in single cell studies?

The CellANOVA model contains explicit terms for sample-specific batch effects, namely **C**^(*i*)^**B**^(*i*)^**V**^⊤^ for the *i*th sample (see Materials & Methods). This allows us to quantify the unwanted variation for each gene in each cell, thus enabling the interrogation of how individual genes are affected by batch in a cell specific manner. We start by visualizing the estimated batch effect terms of the Immunoterapy trial, type 1 diabetes, and mouse radiation datasets through UMAP embedding, which was produced by Harmony-based CellANOVA. See Supplementary Figure 17 for comparable results produced by Seurat-based CellANOVA. As expected, since each sample is a separate batch, the samples are well-separated from one another on the UMAPs. Importantly, we see that the major cell types for each dataset are also separated, to varying degrees, within each sample, indicating that batch effects are highly cell type specific.

Many genes are not only strongly affected by batch, but the magnitude of its batch effect can vary significantly across cells. At the per-gene level, what underlies the cross-cell variation of its batch effect term? To start, we consider the contribution of library size, which is well appreciated to be a technical confounding factor in single cell studies. It is worth noting that we have already normalized by library size during the preprocessing step as we divided each cell’s raw UMI counts by their sum (i.e., we normalized by total counts per cell). Is this simple normalization enough? For each gene, we calculated Pearson’s correlation between its estimated batch effect term (columns of **C**^(*i*)^**B**^(*i*)^**V**^⊤^) and the cell library size within each cell type. As is shown in Supplementary Figure 18, the correlations between library size and batch effect are, on the whole, low and centered around zero, except for epithelial cells from the mouse radiation dataset. This shows that, in some cell types, the technical dependence on library size can not be completely removed by a simple normalization, but that the residual dependence can be captured and removed by CellANOVA.

We also examined the dependence of batch effects on gene expression magnitude (after library size normalization), through Pearson’s correlation of each gene’s batch effect term and its standardized log-transformed expression (columns of **X**^(*i*)^). The distribution of correlations is plotted in Supplementary Figure 18. We found a minor positive correlation between log-transformed expression and batch effect. This shows that highly expressed genes have larger batch effects, even after log transformation.

Next, we examined the pathways that are most affected by batch in each dataset, and asked if any are shared across stud-ies. Note that the three datasets being compared come from different tissues (pancreas, peripheral blood, and intestine), different laboratories, and two different species (mouse and human). CellANOVA estimates the batch-basis matrix **V**, each column of which can be interpreted as a latent “concept” describing the unwanted variation. Each gene has a loading for each latent concept, and the importance of each concept is recorded by its corresponding singular value. We focused on the *k* most important batch-associated concepts (*k* = 5), and computed a weighted-sum of squared loadings across these concepts for each gene, weighted by the corresponding singular values. Intuitively, this weighted sum measures the susceptibility of each gene to batch effects, and thus we call it the batch-susceptibility score (BSS). For each study, we identified the batch-susceptible genes by selecting those with the top 30% highest BSS. Then, we employed a hypergeometric test to discover the enrichment of a priori defined pathways in this batch-susceptible gene set, referring to Molecular Signature Database for pathway information. Figure 7b shows the top ten batch-susceptible pathways of each study. Despite differences in tissue, lab, and species, six out of ten pathways are shared across these three studies: (1) Myc Targets V1; (2) Oxidative Phosphorylation; (3) mTORC1 Signaling; (4) DNA Repair; (5) Myc Targets V2; and (6) Unfolded Protein Response. Supplementary Figure 17 shows that the same pathways could also be found if Seurat, instead of Harmony, is used together with CellANOVA, corroborating these findings.

**Fig. 7:**
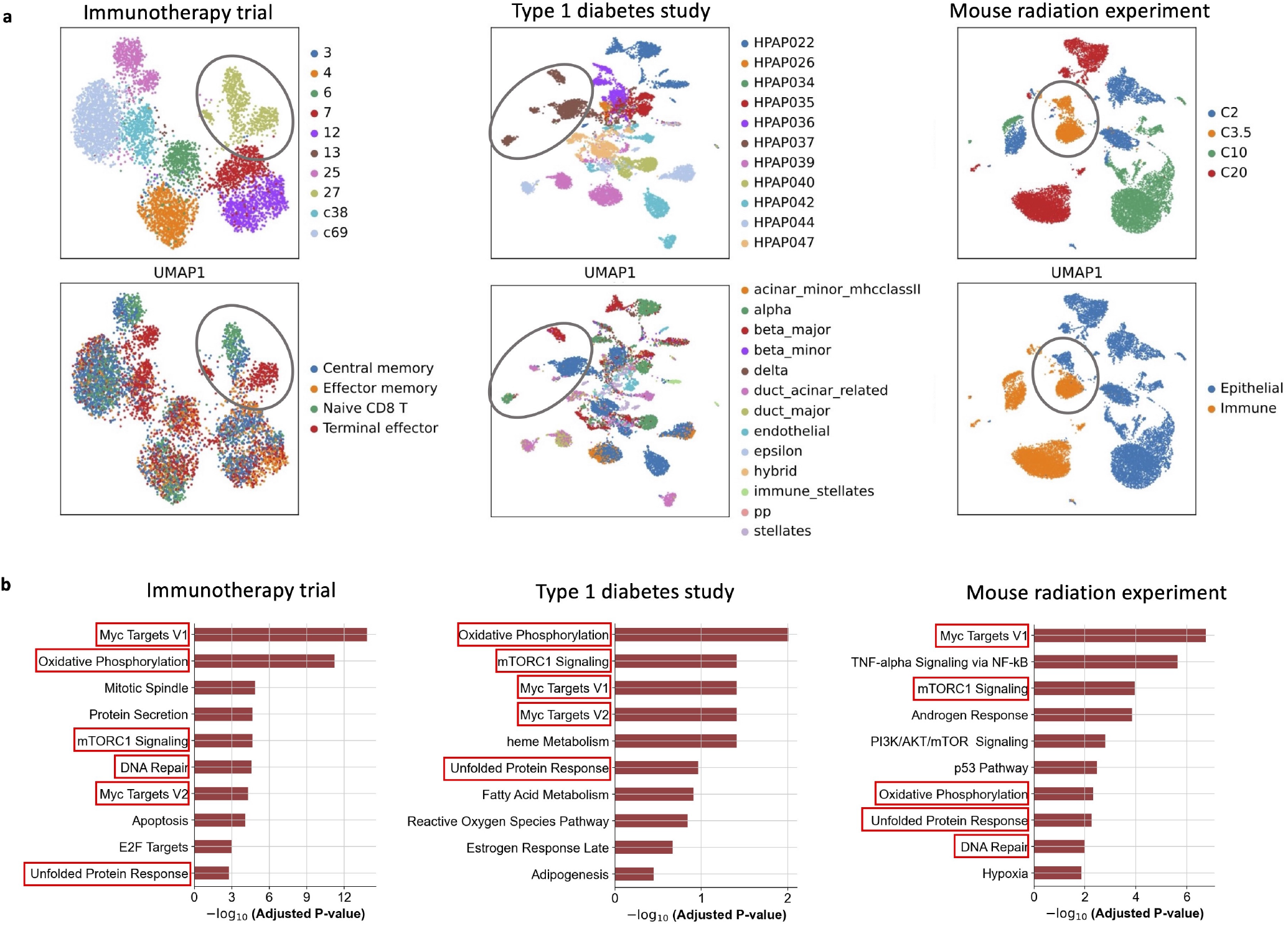
(a) UMAP visualization of batch effects estimated by Harmony-based CellANOVA on three datasets, colored by batch and cell type. (b) Top ten batch-affected pathways of each study based on batch-susceptibility score (BSS) with Harmony-based CellANOVA.

This sharing of batch-susceptible pathways among datasets reflect the fact that, despite differences in tissue and laboratory environment, common technical factors affect single cell sequencing experiments. Subtle variations in tissue processing and handling introduce variations in the level of oxidative and endoplasmic reticulum stress to the cells in different samples, which is why oxydative phosphorylation and unfolded protein response are common sources of high batch variation (37). Oxidative and endoplasmic reticulum (ER) stress lead to coordinated responses: Misfolded proteins can induce reactive oxygen species production, and oxidative stress can disturb the redox environment within the ER thereby further disrupting protein folding (38). Importantly, oxidative stress inhibits mTORC1, an important suppressor of mitochondrial oxidative stress and a key player in cellular stress response and energy metabolism in many cell types (39, 40). Thus, it is not surprising that mTORC1 signalling is a common batch-variable pathway, along with oxidative phospho-rylation and unfolded protein response. The ubiquitous high batch variation of the Myc target genes reflect varying levels of stress-induced cell cycle arrest and cell death across samples (41, 42).

## Discussion

In the analysis of single cell data, integration of cells across samples to remove unwanted variation plays a critical role. Recent advances in the field have brought forth many integration algorithms, each aiming to align cells “belonging to the same state” across multiple samples. However, when the samples are expected to be biologically distinct, there has not been a scientific way to address the question of how aggressively should the cells be aligned. Each integration algorithm has parameter(s) to control the extent of alignment and the resulting uniformity of the samples, but the tuning of such parameters has been left to guesswork. While the stated goal of integration is to remove “batch effect”, batch effects have been challenging to explicitly quantify in the single cell context, and there were no explicit guidelines as to when unwanted batch variation can be separated from biologically meaningful variation.

We developed a new model and analysis framework to explicitly quantify batch effects in single cell data in a cell-state and sample specific way, thus allowing the recovery of variation that is orthogonal to batch effects in single cell integration. This model requires the existence of a set (or sets) of control samples. The inclusion of “control” or “baseline” samples is routine in single cell studies, but such samples are currently used only after integration, e.g., as a baseline for quantifying cell composition changes. By using control samples during the integration step, CellANOVA harnesses good experimental design: control samples should be included not only as biological baselines for comparison but also as representations of the range and diversity of unwanted variation in the experiment. Careful construction of control-pools allows more complete batch effect removal and more sensitive and trustworthy recovery of biological signals.

Through comprehensive benchmarks, we showed that when CellANOVA is applied in conjunction with existing state-of-the-art integration methods, it accomplishes three objectives. First, CellANOVA corrects data distortion introduced by integration, in that it removes batch effects while maintaining maximum similarity to the original data matrix. Second, CellANOVA recovers valid p-values for cross-cell type comparisons, in that it corrects the artificial inflation of cross-cell type differences introduced by current integration methods. Third, CellANOVA allows for the recovery of subtle cell-state-specific differences between samples that were erased during integration. This was shown using both a priori knowledge (in the form of condition labels of the samples) and validation by flow cytometry and by replication in data from an independent study. In our analyses, we applied CellANOVA on initial integrations computed using Harmony and Seurat, but users can choose any integration method. It is important that the initial integration gives a good mixing of the batches, even if that incurs a heavy loss of signal. This is because CellANOVA can recover the biological signal that is lost, but usually can not remove batch effects that are persist after the initial integration.

The CellANOVA model also gives us explicit intuitions on what comprises batch effects in single cell data, and what types of biological signals can be recovered from an integration. Through the batch susceptibility score, we found that a set of shared core pathways have the highest susceptibility to batch effects across data from different labs, tissues, and species. Only the component of biological signals that are orthogonal to the batch latent space can be recovered, and thus we expect that variation in these pathways, if they were not already preserved in the original integration and thus hard-coded in the cell-state encoding, to be refractory to CellANOVA signal recovery.

CellANOVA is a lightweight algorithm that adds only a few minutes to current integration pipelines. The benchmarking procedures we employed in Figure 3 and Figure 4 can be performed on any dataset. They have also been implemented in the CellANOVA package and we believe that they should be routinely used for visualization and diagnostics. A common question should be “do we have a big enough control-pool”. This can be answered by doing the hold-out experiment on the current control-pool, as described in Figure 3a, to see if the hold-out sample is sufficiently well integrated with the remaining control-pool samples. If not, more control samples should be collected to get a more complete representation of the unwanted variation in the data.

## Materials & Methods

### Data preprocessing

All scRNA-seq datasets are transformed as follows before integration: Let 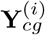 be the raw count for gene *g* in cell *c* in sample *i*. We define 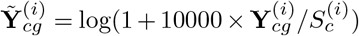, where 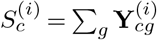 is the library size of cell *c* in sample *i*. Then, the data is centered for each gene across all cells to ob-tain 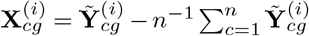. Then, 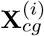 is the value in the CellANOVA model Eq. (1). This is also the starting value for the integration by Seurat, Harmony, and Symphony. For Liger, we followed the pipeline suggested in the software tutorial (https://github.com/welch-lab/liger) (33) and performed library size normalization, highly variable gene selection (*p* = 3000), and scaling without centering.

### Identifiability constraints of CellANOVA model

The model in Eq. (1) is non-identifiable unless some additional constraint is imposed. To ensure identifiability, we make the following assumptions:

i. **V, W**, and 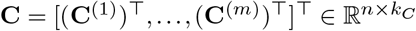 has orthonormal columns where 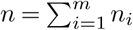;
ii. 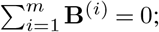
iii. 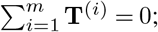
iv. **V**^⊤^**W** = **0**.

Given cell state **C**^(*i*)^, the terms in brackets in Eq. (1) were inspired by the (two-way) ANOVA model (43, 44) where technical variation due to batch effect and biological variation due to treatment condition explains cell-state-specific variations in an additive way.

### Details of model fitting

We fit the model in Eq. (1) by successively carrying out the following three steps.

#### Estimation of cell states

In the first step, we estimate cell states across all *m* datasets. To this end, let

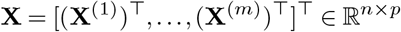

be the stacked data matrix. For a user-selected number of principal components (PCs) *k*_*C*_ *>* 0, we apply Harmony (13) on **X** with *k*_*C*_ PCs to align across dataset labels and use the *k*_*C*_ leading left singular vectors of the Harmony output, collected as the columns of

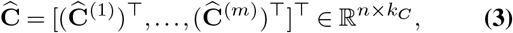

as our estimator of the cell state coding matrix **C** . We could replace Harmony with other comparable batch effect correction methods, such as Seurat (32).

### Estimation of batch effects and main effects

For *i* =1, …, *m*, we regress **X**^(*i*)^ on 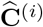 to obtain regression co-efficient matrix **R**^(*i*)^ via ordinary least squares (OLS). Here and after, for two full rank matrices **A***∈* ℝ^*ℓ×q*^ and **B***∈* ℝ^*ℓ×s*^ with *ℓ > s > q*, the OLS regression coefficient matrix from regressing **A** on **B** is given by (**B**^*T*^**B**)^*−*1^**A***∈* ℝ^*s×q*^ .

Now define the within-control average effect

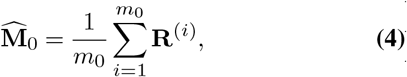

which is the average of regression coefficient matrices in the control/baseline datasets. Define

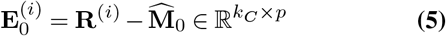

for *i ∈* [*m*_0_] and

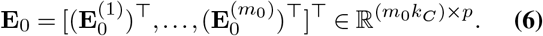

With a user-defined positive integer *k*_*B*_, we estimate **V** by the *k*_*B*_ leading right singular vectors of **E**_0_, collected as columns of 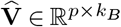 .

Now we estimate the main effect **M** with

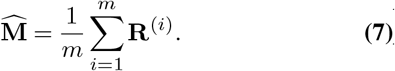

For *i ∈* [*m*], define

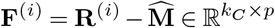

and further define our estimator for **B**^(*i*)^ as

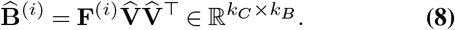

When 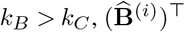 is the unique OLS regression coefficient matrix obtained from regressing 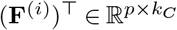 on 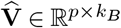 .

### Estimation of treatment effects

For 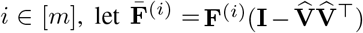 and further define

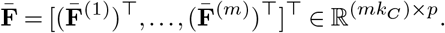

For a user-defined positive integer *k*_*T*_, we estimate **W** in Eq. (1) with 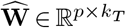 whose columns collect the *k*_*T*_ leading right singular vectors of 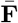 . Furthermore, for each *i*, we estimate **T**^(*i*)^ with

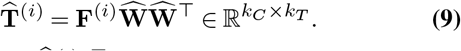

When 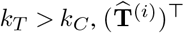 is the unique OLS regression coefficient matrix obtained from regressing 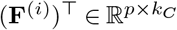 on 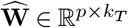 .

By the definitions of 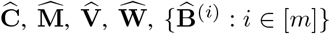, and 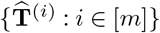, the four identifiability assumptions are satisfied by these estimators.

#### Batch-effect-corrected datasets for exploratory data analysis

In many exploratory scenarios, one may only be interested in removing batch effects while preserving as many biological signals as possible. To this end, the idiosyncratic term **Z**^(*i*)^ may contain valuable signal of interest. When this is the case, the only undesirable term in Eq. (1) is **B**^(*i*)^**V**^T^. To this end, the formula Eq. (2) gives the batch-effect-corrected version of the ith dataset. Effectively, for each dataset, after adjustment with respect to its cell state compositions, we project its difference from the global mean onto the orthogonal complement of the subspace spanned by columns of the estimated batch basis matrix. The sum of the batch-effect-corrected difference and the cell-state adjusted global mean gives the batch-effect-corrected dataset that can be treated as a raw dataset in downstream analysis, such as DEG analysis and gene set enrichment analysis.

### Extension of the basic CellANOVA model to multiple control-pools

Sometimes the desired control/baseline group may consist of datasets collected under multiple conditions. For example, it may contain scRNA-seq data collected on a number of healthy controls and on all diseased subjects prior to treatment. In this case, assume there are *q* disjoint groups of controls, denoted by 𝒞_1_, …, 𝒞_*q*_, under different conditions such that the union 𝒞_1_ *∪… ∪*𝒞_*q*_ = [*m*_0_] covers all control datasets. For any set 𝒞, let |𝒞| denote its cardinality. For *j* = 1, …, *q*, define 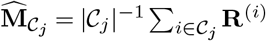 . Let **1**_*E*_ denote the indica-tor of an event *E*. For each *i ∈* [*m*_0_], replace the definition of 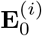 in Eq. (5) with

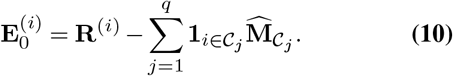

In other words, after cell state composition adjustment, we replace the contrast of a control dataset against the mean over all control Eq. (4) with the contrast of it against its group mean, as we do not want to contaminate estimated batch effects with differences among means of different control groups. Then, we define **E**_0_ as in Eq. (6) with the above new definition of each 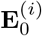 . Finally, we estimate **V** with 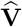 which collects the leading *k*_*B*_ right singular vectors of **E**_0_ as its columns.

### Benchmarking the effectiveness of batch effect removal and distortion correction (Figure 3)

#### Details of methods execution for hold-out analysis

For this analysis, we focused on the control-pool samples within each dataset and employed a hold-out strategy for methods benchmarking. In each experimental run, we designated one control sample as a pseudo-treatment sample (holdout set) and used the remaining control samples as the pseudo-control-pool for CellANOVA. To ensure comparability, the quality control and low-quality-cell removal steps were standardized across all methods for each dataset. We ran the suggested workflow of each method to perform data integration. For CellANOVA, we only used designated control samples to estimate the batch variation basis **V** in the second step of model fitting, while all samples (including the held out pseudo-treatment sample) were used to estimate the main effect **M**, treatment-effect variation basis **W**, and cell states **C**. For Harmony, we integrated all samples together using the `harmony_integrate` function in the Python package `Scanpy` (v1.8.1) with default parameters, ignoring the treatment-control design. For Seurat V4, we followed the reference-based integration workflow, specifying the samples in the training set as the reference and the pseudo-treatment sample as the query. The `FindIntegrationAnchors` function with `reduction = “rpca”` and the `IntegrateData` function from the R package `Seurat` (v4.3.0) were used to integrate pseudo-control-pool samples and the pseudo-treatment sample. For Symphony, we set the training control samples as the Symphony reference and the fake treatment sample as the query object. Following the suggested pipeline, we used the `buildReferenceFromHarmonyObj` and `mapQuery` functions from the R package `symphony` (v0.1.1) with default parameters to construct the reference and to integrate the query with the reference, respectively. For Liger, we followed their workflow for integrating multiple single-cell RNA-seq datasets and used the `optimizeALS` and `quantile_norm` functions from the R package `rliger` (v1.0.0) with default parameters to perform joint matrix factorization and quantile normalization.

### Evaluation metrics

We employed the following evaluation metrics in Figure 3:

1. iLISI. To assess local batch mixing of the integrated gene expression data, we used LISI integration (iL-ISI) proposed by Korsunsky et al. (13). It measures the effective number of batches in the neighborhood of a cell. Higher iLISI values indicate better mixing of cells from different samples or batches in the integrated space. We used function `pca` in Python package `Scanpy` (v1.8.1) to perform principal component analysis on the batch-corrected data and then used function `compute_lisi` in Python package `harmonypy` (v0.0.6) to compute iLISI scores. For comparison, all methods in benchmarking utilized the first 15 components to compute iLISI, with all other parameters at their default values.
2. Gene expression correlation. To assess the severity of global distortion, we computed the Pearson’s correlation coefficient between each cell’s gene expression vector before and after correction. A higher correlation indicates milder global distortion. The function `corrcoef` in Python package `NumPy` (v1.20.3) was used to compute Pearson’s correlation.
3. Predicted p-value for differential expression gene test. To evaluate gene signal distortion in the batch correction process, we employed a p-value comparison method inspired by the train-test-split concept commonly used in statistics and machine learning. In step 1, we divided samples in the control-pool into two sets: a training set used for model fitting, and a test set for evaluating model performance. Without loss of generality, let 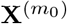 denote the held-out testing sample, and **X**^(1)^,…, 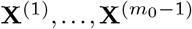 represent the remaining *m*_0_*−*1 training samples. In step 2, using training samples, we followed the CellANOVA pipeline and fitted the model. Specifically, we estimated cell state ma-trices 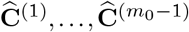, main effect 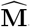, and batch-induced modes of expression variations 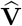. In step 3, we estimated the cell state matrix 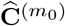 for the held-out control sample by applying Symphony mapping. To achieve this, we used `mapQuery` function in the R package `symphony` (v0.1.1) with default parameters, setting the held-out sample 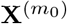 as the query object, and the other samples 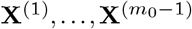 as the reference object. In step 4, we predicted the batch-corrected data for the held-out sample with: 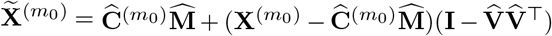. In step 5, we performed differential expression analysis across cell types using 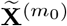 and computed multiple-testing adjusted *P* -value for each gene. We also used uncorrected held-out sample 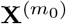 and performed the same differential expression analysis, again obtaining adjusted p-values for each gene. Wilcoxon signed-rank test was used for DE analysis and Benjamini-Hochberg procedure was used to control the false discovery rate. Note that the pre- and post-correction p-values are computed using exactly the same sets of cells, all derived from the same sample. Since they all come from the same sample, the differences between these cells are not influenced by batch effects, and thus, ideally the post-correction p-values should resemble their pre-correction counterparts. We compared the pre- and post-correction adjusted p-values, with a high correlation indicating minimal gene-level distortion.

### Treatment effect detection and estimation (Figure 4)

#### Details of methods execution

In the type 1 diabetes (T1D) dataset, the 11 healthy individuals served as the control-pool. Similarly, for the immunotherapy trial dataset, the 10 samples collected at baseline (time 0) before treatment served as the control-pool. In the mice radiation data, the Sham-irradiated control mice were designated as the control-pool. Samples from the control-pool were used to estimate the batch variation basis **V** for CellANOVA, and to build the reference map in Seurat V4 and Symphony. Harmony and Liger integration were performed across all samples together, ignoring the control-treatment design. The detailed data integration procedures for each method were the same as those used in the previous section. After integration, we recovered batch-corrected gene expression measurements for each cell. For CellANOVA, we first extracted the main and treatment effects from the model, and then combined them 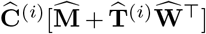 to detect biological signals and evaluate performance. Harmony, Seurat V4, and Symphony output the cell-specific batch-corrected embeddings and a gene-loading matrix. We recovered batch-corrected measurements for each gene in each cell by multiplying batch-corrected embeddings with the gene-loading matrix. Liger identified a set of shared- and dataset-specific latent factors (meta-genes) that corresponded to biological or technical signals, and calculated meta-gene expression for each cell. We recovered batch-corrected gene expression by multiplying meta-gene expression with the meta-gene loading.

#### Evaluation metrics

We employed the following evaluation metrics in Figure 4:

1. Out-of-sample nearest-neighbor proportion. This metric is used to evaluate the extent of preservation of meaningful biological variation across samples. In the first step, we performed principal component analysis on the batch-corrected data and selected the first *n*_*pc*_ components as the features for the k-nearest neighbors algorithm. We set *n*_*pc*_ to 20 for all methods. In the second step, for each cell, we identified its nearest neighbors among cells from the other batches (that is, if the cell comes from sample *i*, we exclude all cells from sample *i* in the nearest neighbor search). We used function `NearestNeighbors` from Python package `scikit-learn` (v1.0.2) with default parameters, using those out-of-batch cells as the training set, then the `kneighbors` function from the same package to predict the *k* (*k* = 30) nearest neighbors in the training set for each cell. In the third step, we computed the proportions of these out-of-batch nearest-neighbor cells belonging to each treatment group. R function `geom_density` from package `ggplot2` (v3.4.1) with smoothing bandwidth `bw=0.05` was used to generate the kernel density plots.
2. Differential gene expression analysis (DEG). To perform differential gene expression analysis between cell types or between conditions, we used Wilcox signed-rank test, implemented in the function `wilcox.test` from the R `Stats` (v4.2.2) Package. To adjust p-values for multiple comparisons, Benjamini & Hochberg procedure was applied using the function `p.adjust`.
3. AUC and ROC. Area Under the Curve (AUC) of the Receiver Operating Characteristic (ROC) curve was used to assess the performance of the marker gene prediction task based on the batch-corrected data produced by different integration methods. For each gene, we performed DEG with batch-corrected data across conditions and assigned an adjusted *P* -value (as described above), which was used for marker gene prediction. We used marker vs. non-marker in the simulation dataset as the ground truth. The number of genes that were detected as markers and are true differentially expressed genes (DEGs) is denoted as true positives (TP). The number of genes detected as markers but not true DEGs is referred to as false positives (FP). True positive rate is the proportion of TP among all positive detections. False positive rate is the proportion of FP among all positive detections. Functions `geom_roc` and `calc_auc` in R package `plotROC` (v2.3.0) were used to plot ROC curves and compute AUC values, respectively.
4. Gene Set Enrichment Analysis (GSEA) (45, 46). To identify biologically relevant gene sets associated with different cell groups, such as cell types or time points, thereby assessing signal preservation after batch correction, we performed Gene Set Enrichment Analysis. In the first step, we conducted a t-test for each gene using batch-corrected data between two cell groups (such as two cell types or two time points). Then, we ranked genes using *t* statistics. Next, we followed the standard protocol outlined in the tutorial of the Python package `GSEApy` (v1.0.4), with the ranked gene list as input and all parameters set to default. To mitigate any bias caused by uneven distribution of cells among batches, we subsampled the data to ensure that each batch contained no more than *n* cells, with *n* set to 300. The main function we used is `prerank` from package `GSEApy` (v1.0.4), which employs permutation tests to determine whether a priori defined sets of genes show statistically significant enrichment at either end of the ranking. MSigDB Hallmark 2020 database was used as the reference gene sets. GSEA plots were generated using `gseaplot` function from the same Python package.

### Details of analysis of spatial transcriptomics data for Figure 6d, 6e, and Supplementary Figure 14

Deconvolution for all samples was performed using RCTD using the MULTI setting with the maximum number of cell types per spot equal to 4. In each spot, cell types that were reported to comprise less than 5% of the spot were filtered out and the cell type proportions were renormalized. Gene sets corresponding to specific pathways were pulled the hallmark gene sets from GSEA | MSigDB | Browse Human Gene Sets (gsea-msigdb.org). For each gene set and Visium sample, spot-specific pathway enrichment scores were computed via the ‘AddModuleScore’ function in Seurat.

### Details of trajectory and pathway activity analysis for Figure 6f

After preprocessing, we first conduct Harmony integration and CellANOVA signal recovery on proximal tubule cells (PT), including three subclusters (PT_S1, PT_S2, PT_S3), as well as injured PT cells (iPT), utilizing both scRNA-seq and snRNA-seq data. Next, we performed Leiden clustering and computed the PAGA graph introduced by (47). Then, we recomputed the embedding using PAGA-initialization. We selected PT cell as the root for the computation of diffusion pseudotime (48). In order to calculate TNF-alpha signaling via NF-kB pathway activity for cells along the trajectory, we adopted the methodology described in (49). The gene sets used in this analysis were downloaded from (https://www.gsea-msigdb.org/gsea/msigdb/human/geneset/HALLMARK_TNFA_SIGNALING_VIA_NFKB.html). The main functions for the above analysis are from Python package `Scanpy` (v1.8.1): `tl.paga`, `tl.draw_graph`, `tl.dpt`, `tl.score_genes`.

### Details of simulation model for Supplementary Figure 15

A negative binomial distribution was used to generate gene counts *y*_*cg*_ based on a gene-and-cell specific mean *μ*_*cg*_ and a fixed dispersion parameter *θ* = 0.35. For cell types CT1-CT5, the distribution of the expression mean *μ*_*cg*_ was the same across control and treatment batches. Specifically, the distribution of means of marker genes was Uniform(2.5, 3.5) and the distribution of means of other background genes was Uniform(1, 2). For cell type CT6, the distribution of marker gene means in the five control batches was the same as above, while that of marker gene means in two treatment batches was scaled up by an additive factor of 1.5 resulting in *μ*_*cg*_ ∼ Uniform(4, 5). Batch effects were added as a small shift to the gene expression means where the shift were i.i.d. random numbers sampled from a Log-Normal distribution (mean and standard deviation on the log scale to be 0.01 and 0.35, respectively).

### Computation of batch susceptibility score (BSS)

Recall that in the second step of CellANOVA model fitting, we estimate the batch variation basis matrix by performing a singular value decomposition on **E**_0_ (defined in Eq. (6)), which is the regression coefficient matrix in the control/baseline datasets after demeaning. Then the estimated batch variation matrix 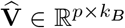 is composed of *k*_*B*_ leading right singular vectors, where the corresponding singular values are denoted as 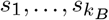 . We define batch-susceptibility score (BSS) for gene *g* as

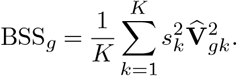

where we set *K* = 5. Intuitively, each column of 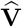 can be interpreted as a latent “concept” describing the unwanted variation. Each gene has a loading for each latent concept, and the importance of each concept is recorded by its corresponding singular value. Batch susceptibility score (BSS), therefore, measures the susceptibility of each gene to batch effects.

### Experiments and datasets

#### Mouse radiation therapy dataset

C57BL/6J mice (Jackson Labs, Bar Harbor, ME) were divided into two experimental groups of sham-irradiated control mice (C) and conventional-dose-rate-irradiated mice (SR). Whole abdominal irradiation with standard PRT (0.9 *±*0.08 Gy/s) was delivered as previously described. At days 2, 3.5, 10, and 20 post-irradiation, intestinal segments of two or more mice from each group were harvested, and single cells were isolated and sequenced from the epithelial and lamina propria layers of the organ. The single cells from the two mice were then pooled in their respective fractions, and flow cytometry was used to enrich for ten-thousand live cells from each fraction but not for any cell populations. Single cell emulsions were obtained using the 10x Chromium Controller, and libraries were prepared using the Chromium Single-Cell 3’ Library & Gel Bead Kit v2 (10x Genomics) following the manufacturer’s protocol. Libraries were sequenced on an Illumina NextSeq using a NextSeq 500/550 v2.5 High Output Kit (Illumina).

#### Immunotherapy trial dataset

The immunotherapy trial dataset was retrieved from Divij et al. (30) In the original study, Divij et al. (30) isolated PBMC cells from specific patients and sorted them into live CD8+ cells using a BD FACs Aria II sorter. The sorted cells were then encapsulated into GEMs using a 10x Chromium Controller and transformed into libraries following the Chromium Next GEM Single Cell 5’ Reagent Kits v2 (Dual Index) Protocol. Subsequently, the libraries were sequenced on a NovaSeq 6000 platform. The obtained sequencing data were processed using the Cell-Ranger pipeline v5 from 10x Genomics, with BCL files being converted into FASTQ format and aligned to the human genome (GRCh38) to produce count matrices. Doublets were then identified with R package `DoubletFinder`. Cell type annotations were generated using function `SingleR` from R package `SingleR` (v1.10.0). A collection of 114 bulk RNA-seq samples of sorted immune cell populations from GSE107011 (50) were used as the reference to label CD8 T cells.

#### Type 1 diabetes study dataset

The scRNA-seq data of the type 1 diabetes study was retrieved from Fasolino et al. (29). Doublets removal was performed by Fasolino et al. (29) using `DoubletFinder`. Cell type annotations were shared by the authors of (29). They utilized the R package `Garnett` for initial cell classification and validated the cell type assignments by integration and label transfer. We randomly sub-sampled 30,000 cells for our study.

#### Human kidney multi-omics atlas dataset

The single-cell RNA-seq, single-nuclei RNA-seq, and VISIUM data we analyzed in Figure 6 are retrieved from Abedini et al. (31). Cell type labels and disease group information are provided by the original study. For independent validation of our findings using Abedini et al.’s data, we also downloaded human kidney single-cell RNA-seq data by Lake et al.(36) from the Kidney Precision Medicine Project (KPMP). We subseted CKD and Healthy donors from data generated by KPMP (https://www.kpmp.org.): DK133081, DK133091, DK133092,DK133093, DK133095, DK1330971, DK114866, DK114908, DK133090, DK133113,DK133766, DK133768, DK114907, DK114920, DK114923, DK114933, DK114886. Data downloaded on [June 15, 2023].

## Supporting information

No need for link text

## CODE AVAILABILITY

All code used in this study, including the CellANOVA software and the analysis code, can be found at https://github.com/Janezjz/cellanova.

## DATA AVAILABILITY

The mouse radiation therapy dataset from is available by reasonable request upon journal publication. The immunotherapy trial dataset is obtained from Mathew et al. (30) by request. Type 1 diabetes study dataset (29) is available at GEO under accession GSE148073. Human kidney multi-omics atlas dataset (31) is available at GEO under accession GSE211785. Single-cell RNA-seq data by Lake et al. (36) can be downloaded from (https://www.kpmp.org., including DK133081, DK133091, DK133092,DK133093, DK133095, DK1330971, DK114866, DK114908, DK133090, DK133113,DK133766, DK133768, DK114907, DK114920, DK114923, DK114933, DK114886.)

## ACKNOWLEDGEMENTS

This work was funded in part by grants from the National Science Foundation DMS-2210104 (Z.M.), the National Institutes of Health R01-HG006137-11, U2C-CA233285 (N.R.Z.), Mark Foundation Center for Radiobiology and Immunology (N.R.Z., A.M.). D.M. was supported by the Parker Institute for Cancer Immunotherapy.

## AUTHOR CONTRIBUTIONS

Conceptualization: ZZ, ZM, NRZ

Algorithm Development: ZZ, ZM, NRZ

Software implementation: ZZ

Design of computational experiments: ZZ, ZM, NRZ

Immunotherapy trial experiment: DM

Mouse irradiation experiment: CM, TL

Kidney disease experiment: KS

Data Analysis: ZZ (with help from DM, TL, KM, SH, ZM, and NRZ)

Manuscript writing: ZZ, ZM, NRZ with feedback from TL, DM, KM

Supervision: AJM, EJW, KS, ZM, NRZ

