## Supplementary material for "Signal recovery in single cell batch integration": No need for link text

### Supplementary Figures

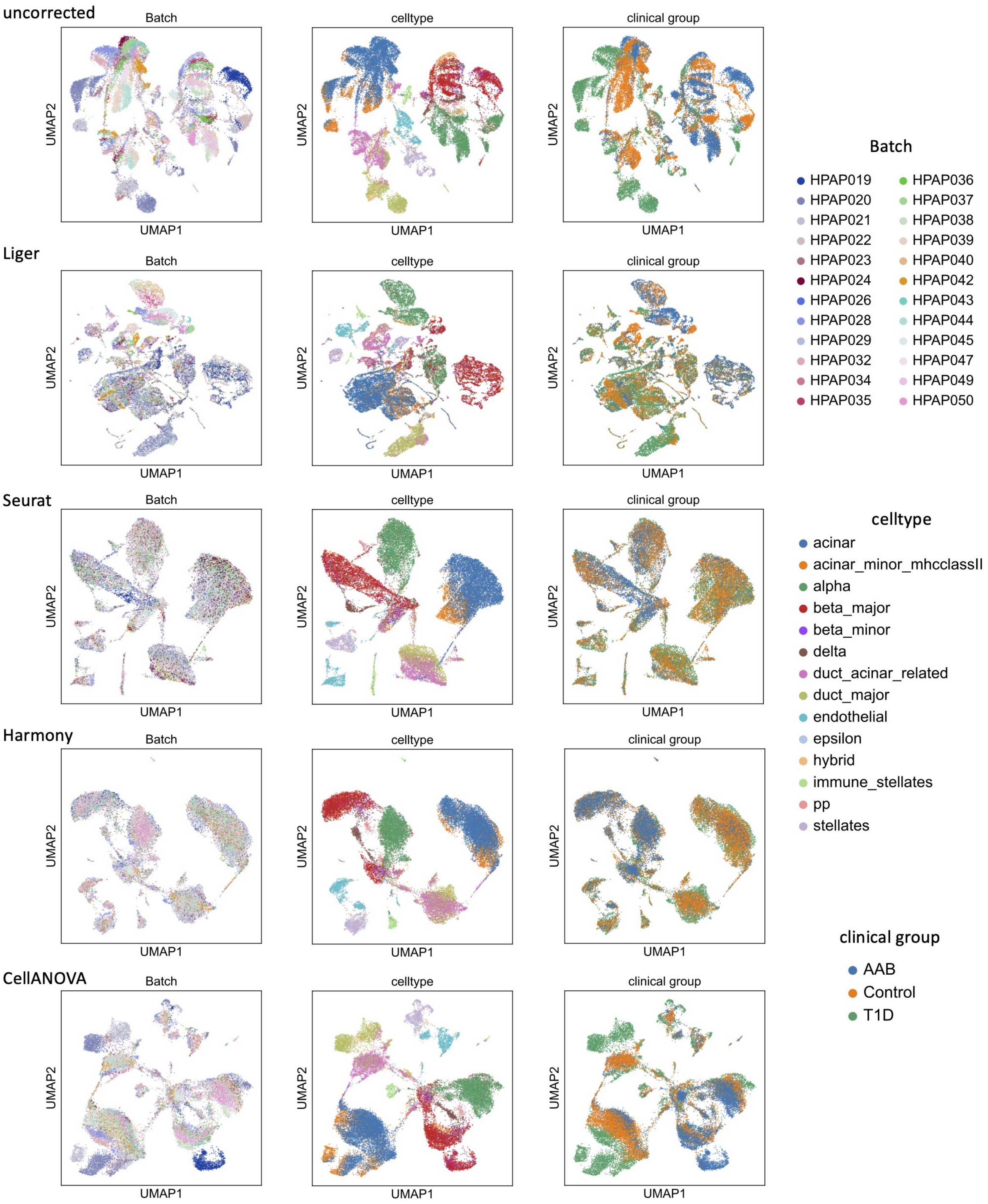


Fig. 1: UMAP visualizations for type 1 diabetes dataset.

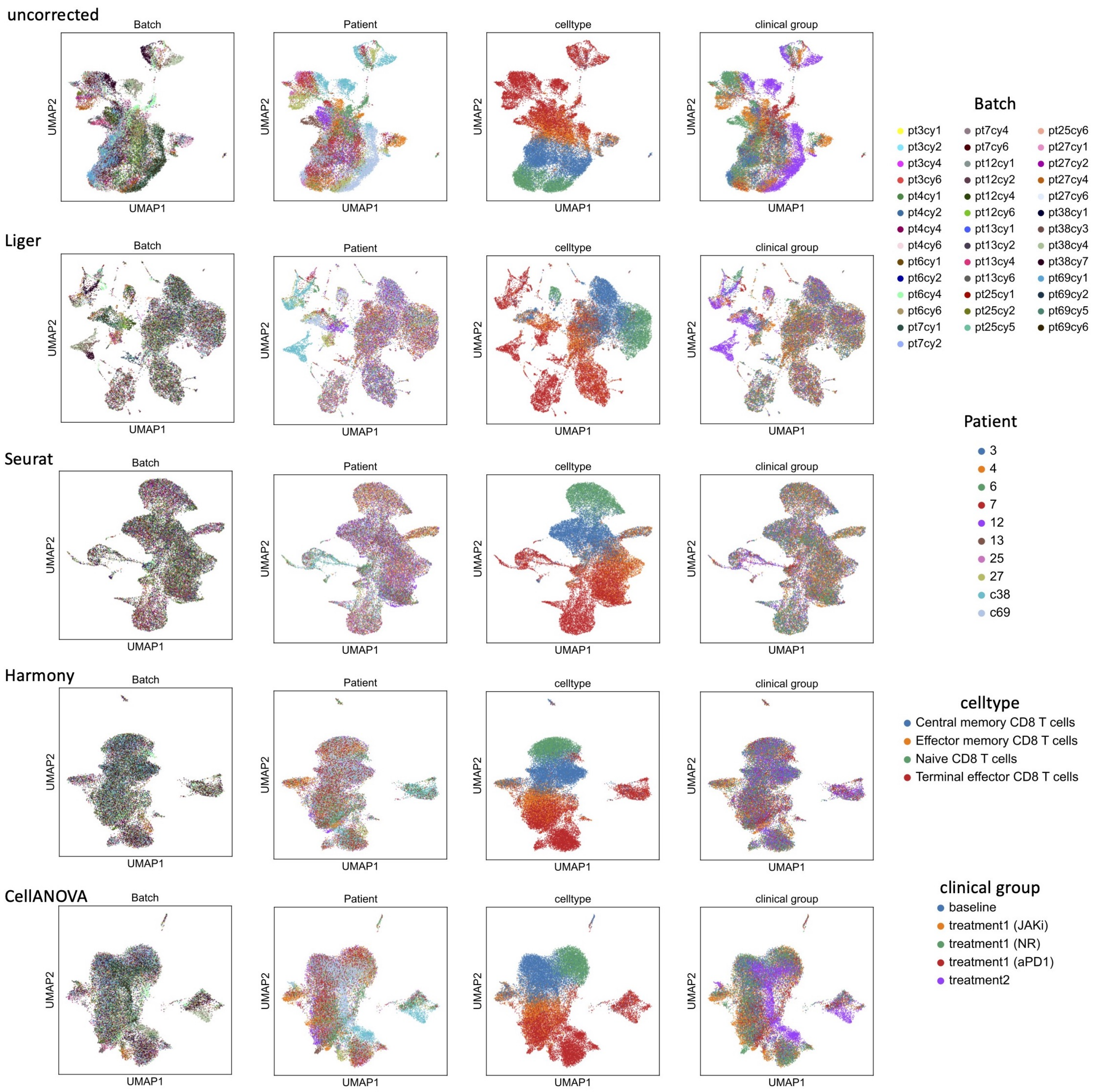


Fig. 2: UMAP visualizations for immunotherapy trial dataset.

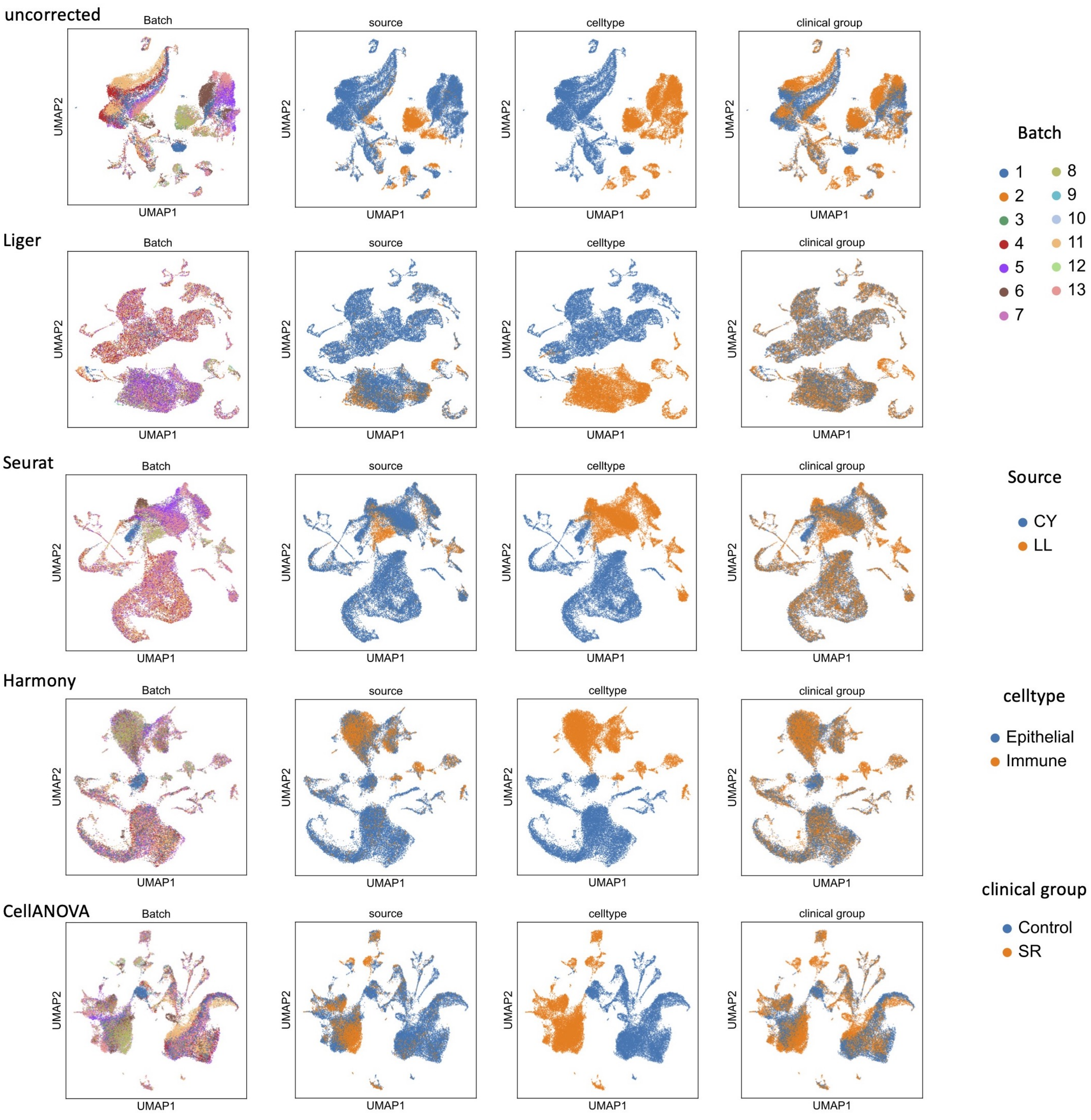


Fig. 3: UMAP visualizations for mouse radiation experiment dataset.

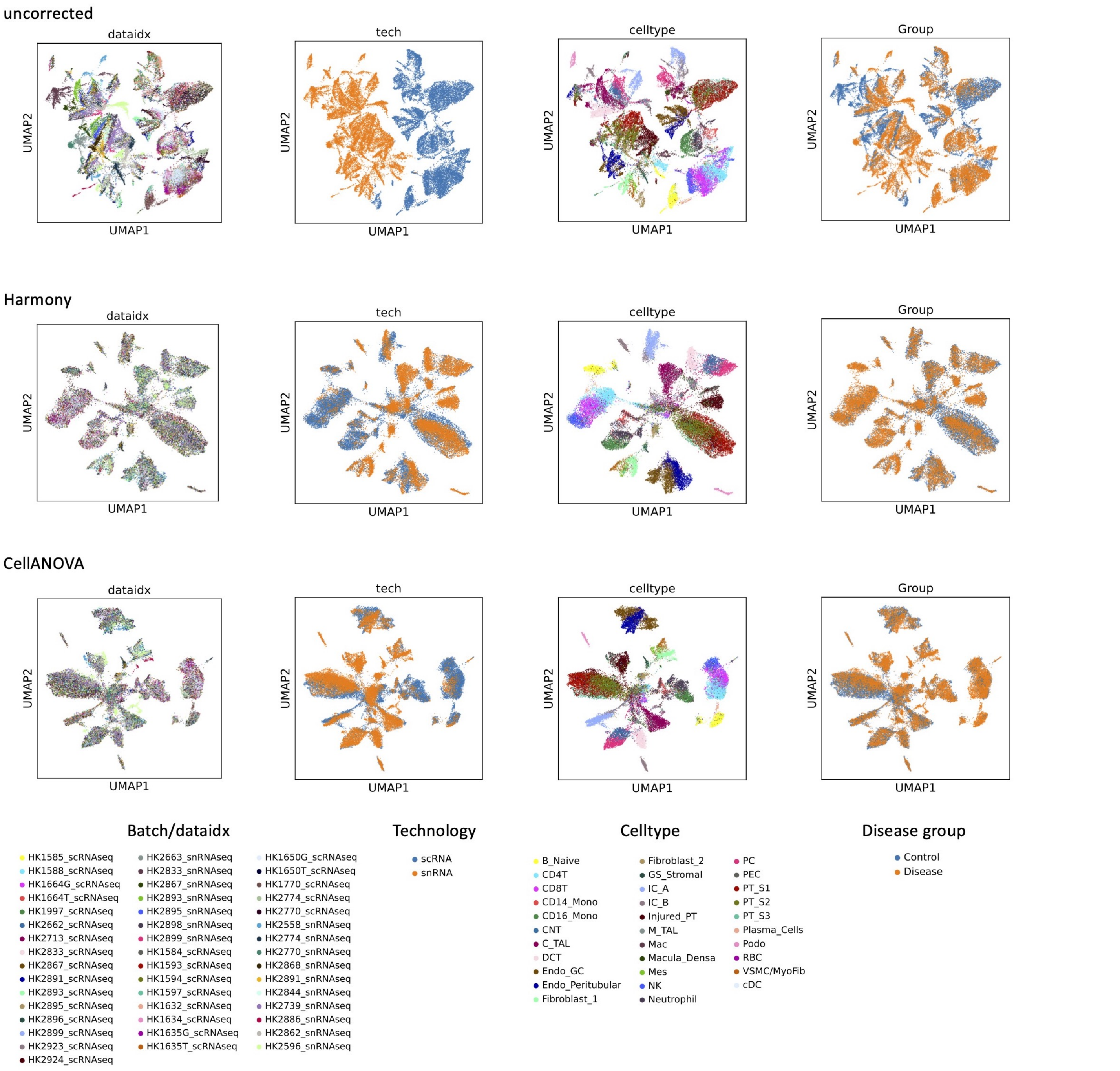


Fig. 4: UMAP visualizations for multi-omics kidney atlas dataset.

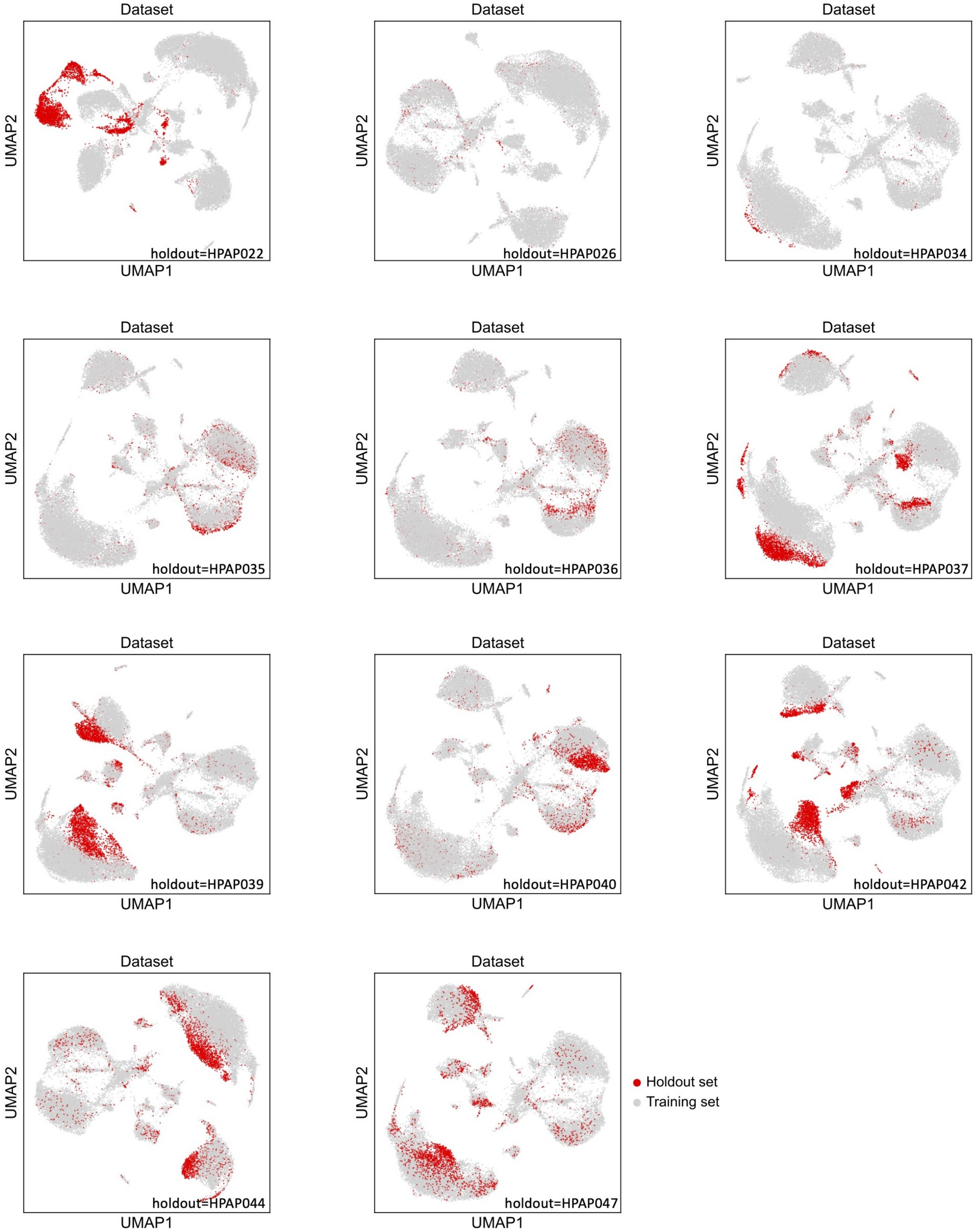


Fig. 5: UMAP visualizations of the leave-one-out experiment performed on the Type 1 diabetes dataset.

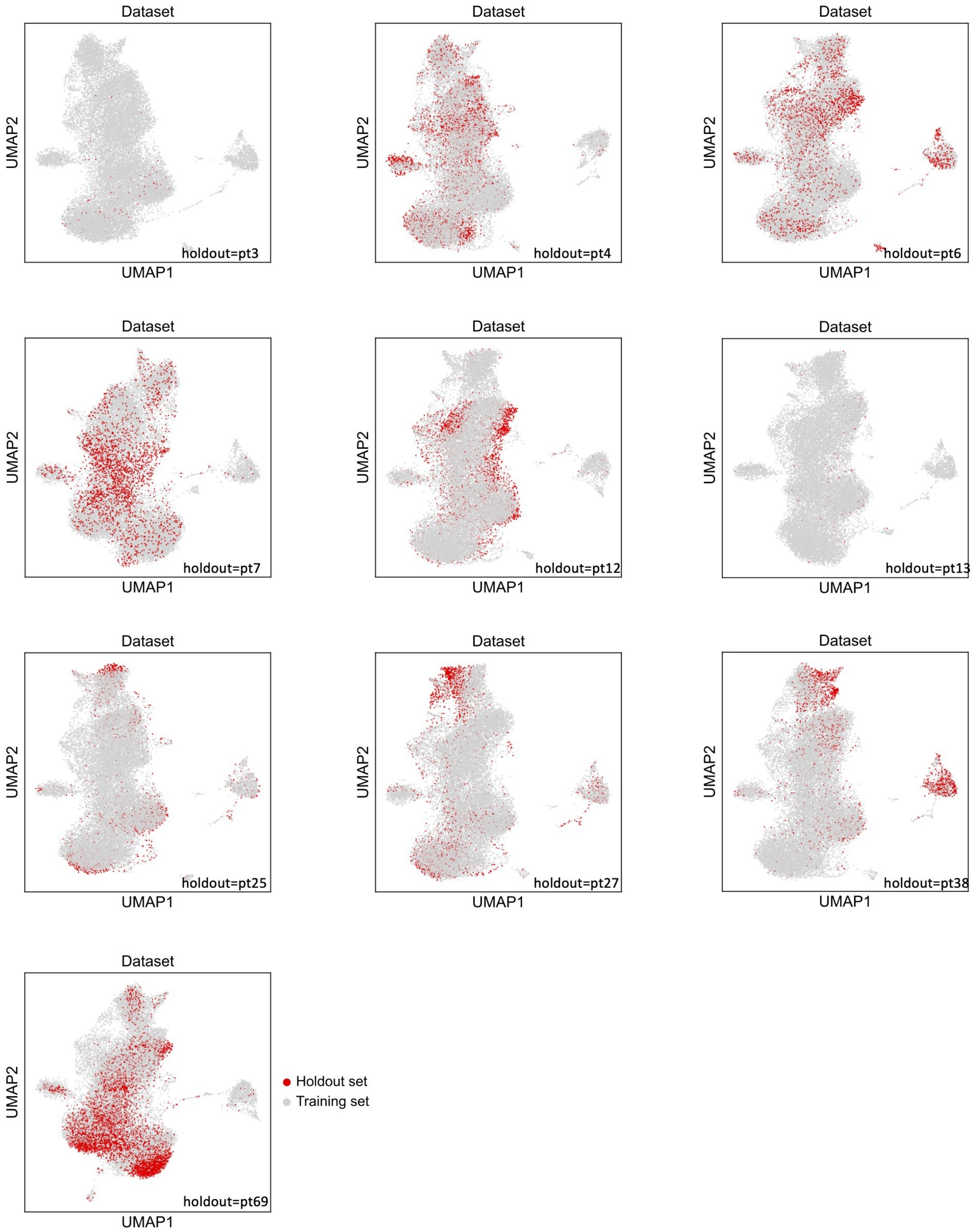


Fig. 6: UMAP visualizations of the leave-one-out experiment performed on the immunotherapy trial dataset.

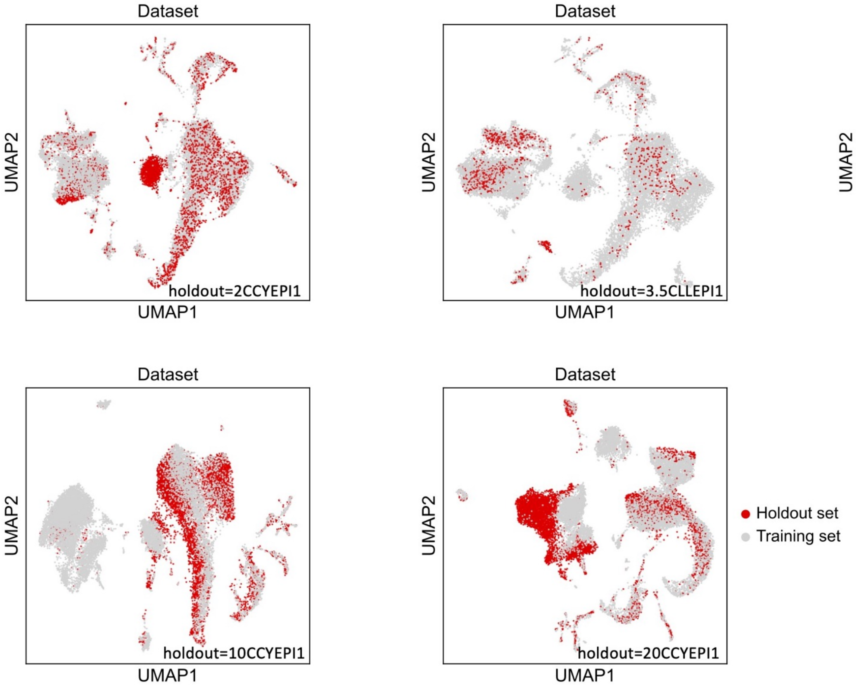


Fig. 7: UMAP visualizations of the leave-one-out experiment performed on mouse radiation experiment dataset.

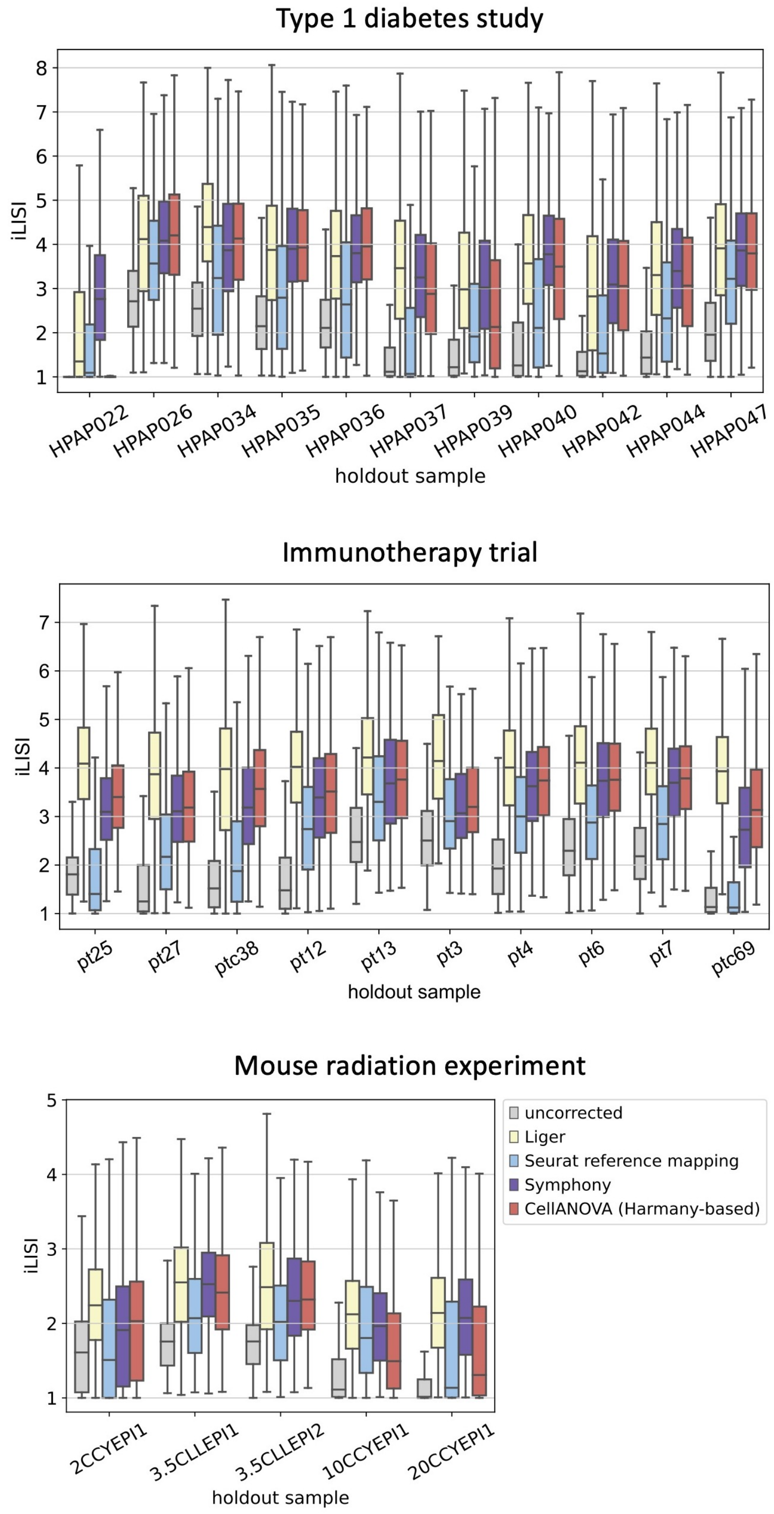


Fig. 8: iLISI scores of the fake treatment sample after batch correction (with different methods) in each hold-out experiment.

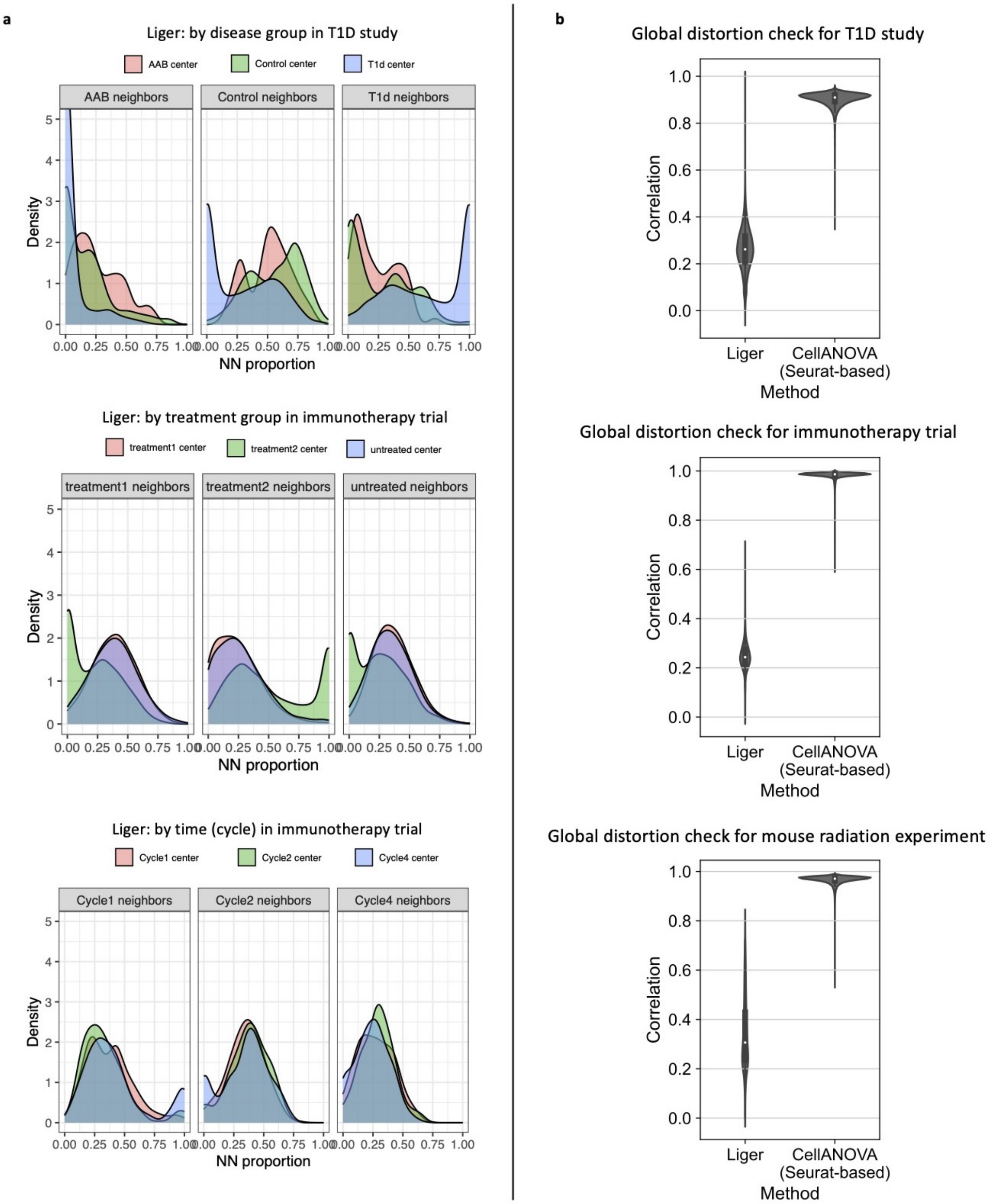


Fig. 9: (a) Out-of-batch nearest-neighbor composition based on Liger-integrated data. (b) Global distortion check for Liger, compared with Seurat-based CellANOVA. Violin plots show the correlations between pre- and post-correction gene expressions per cell.

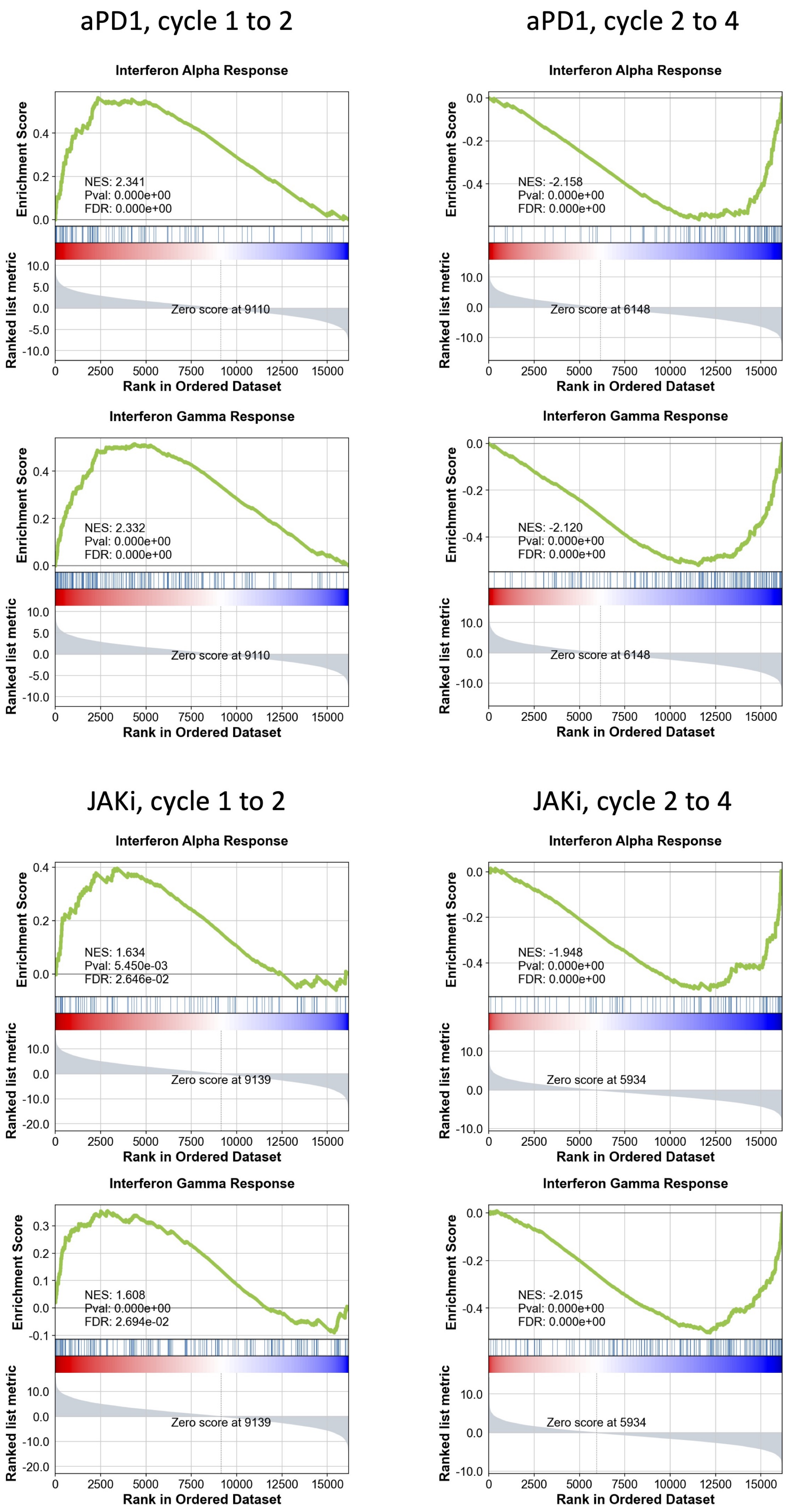


Fig. 10: GSEA plot for IFN-$\alpha$ and IFN-$\gamma$, comparing cycle 1 vs. cycle 2, cycle 2 vs. cycle 4, within aPD1 and JAKi group. A positive normalized enrichment score (NES) from GSEA indicates higher enrichment in the later time points, thereby demonstrating cumulative up-regulation of the set of genes. Conversely, a negative NES score indicates enrichment in the earlier time points and demonstrates cumulative down-regulation of the set of genes in the later time points.

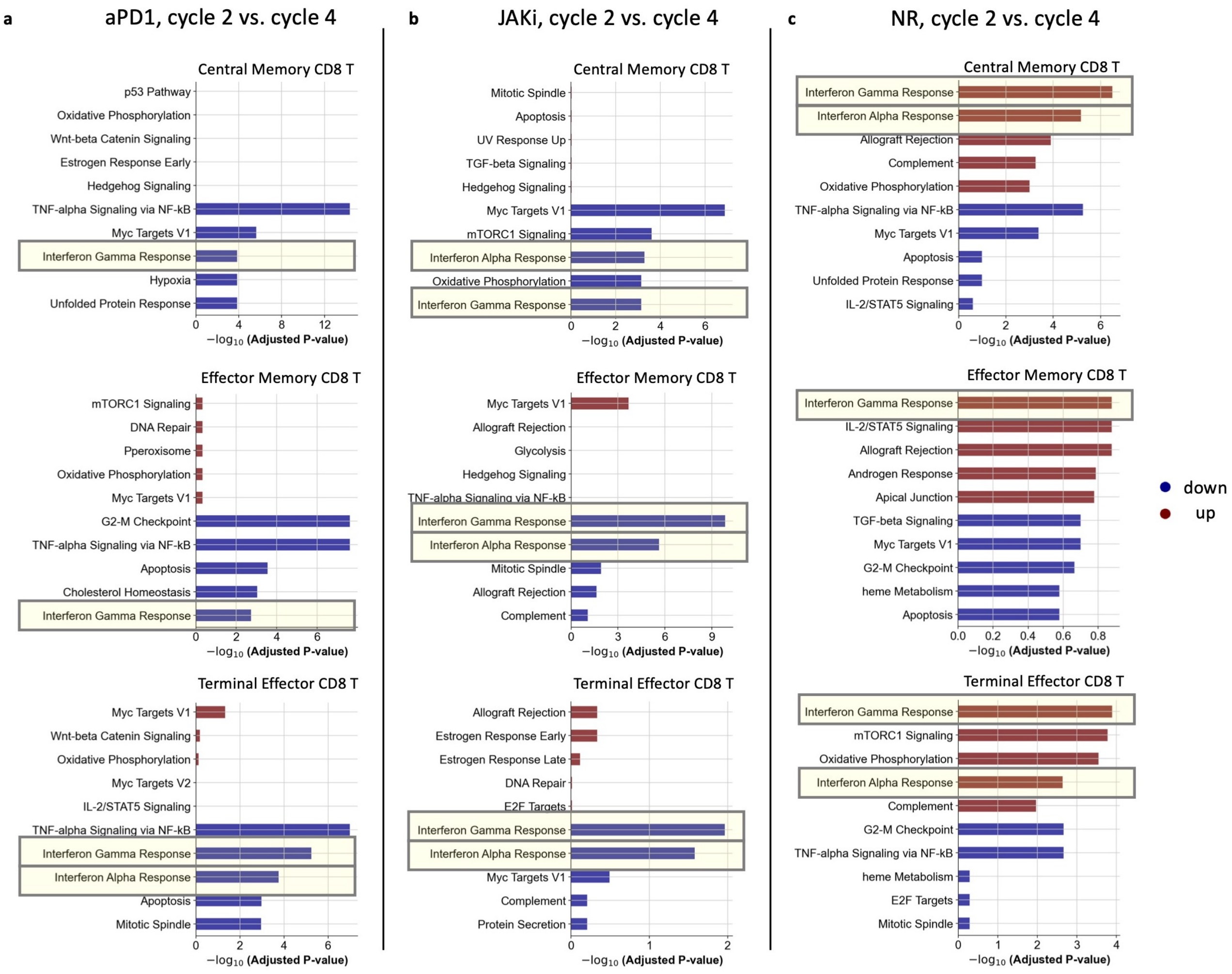


Fig. 11: Cell-subtype-specific gene set analysis within each response group between cycle 2 and cycle 4 after Seurat-based CellANOVA integration. Top 5 up-regulated and down-regulated pathways in cycle 4 compared to cycle 2 are shown.

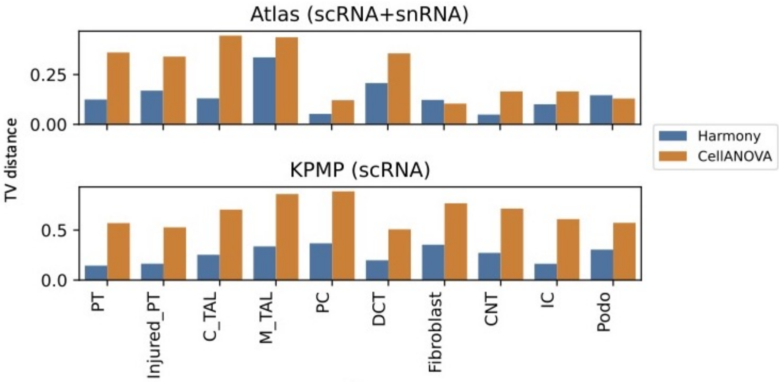


Fig. 12: Total variation distance between two distributions: distribution of out-of-batch nearest neighbor proportion from disease condition around diseased cells; distribution of out-of-batch nearest neighbor proportion from control condition around diseased cells.

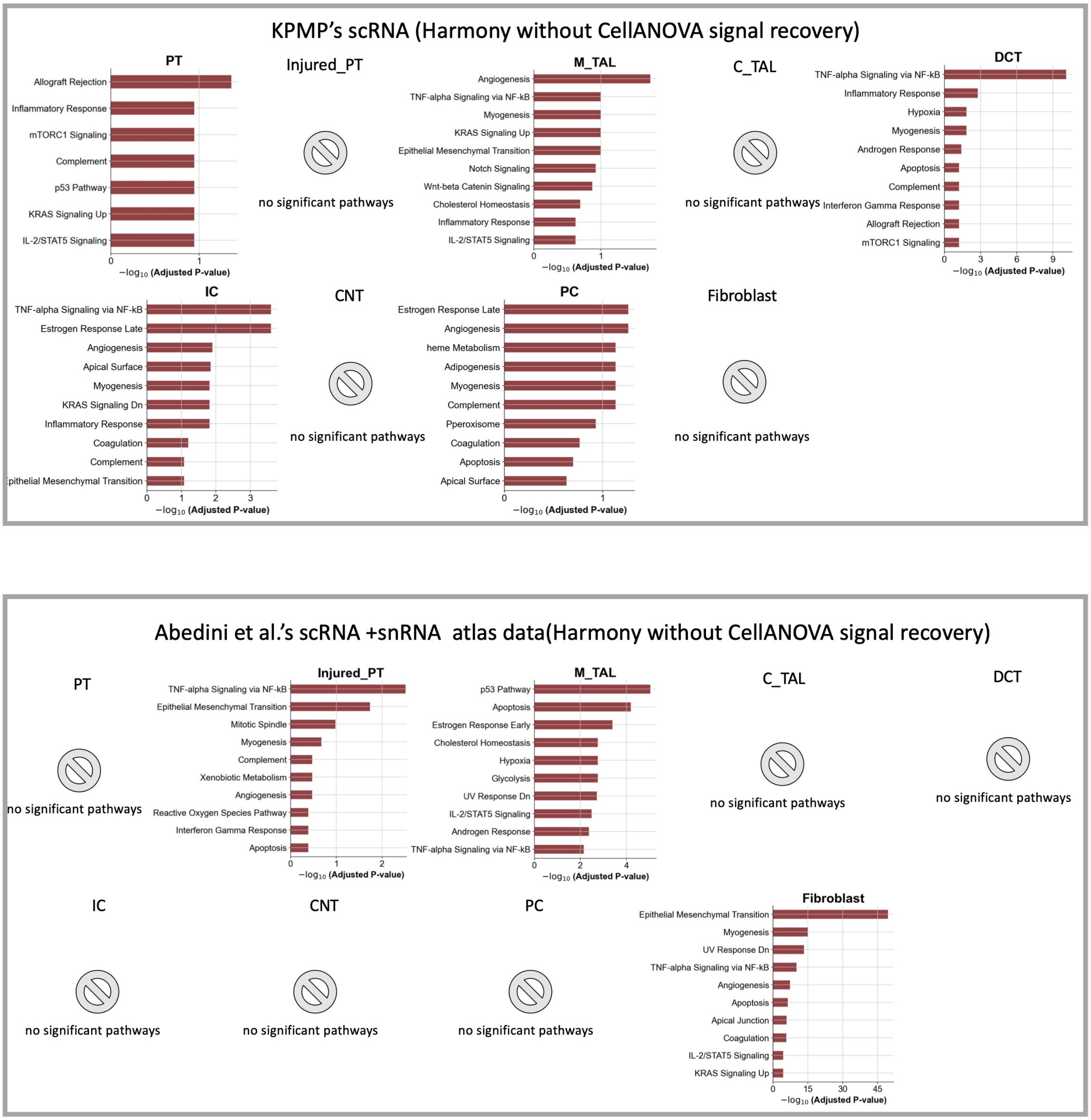


Fig. 13: Top ten upregulated pathways identified within the disease condition for each specific cell type in the Abedini et al. data and the KPMP data without CellANOVA recovery.

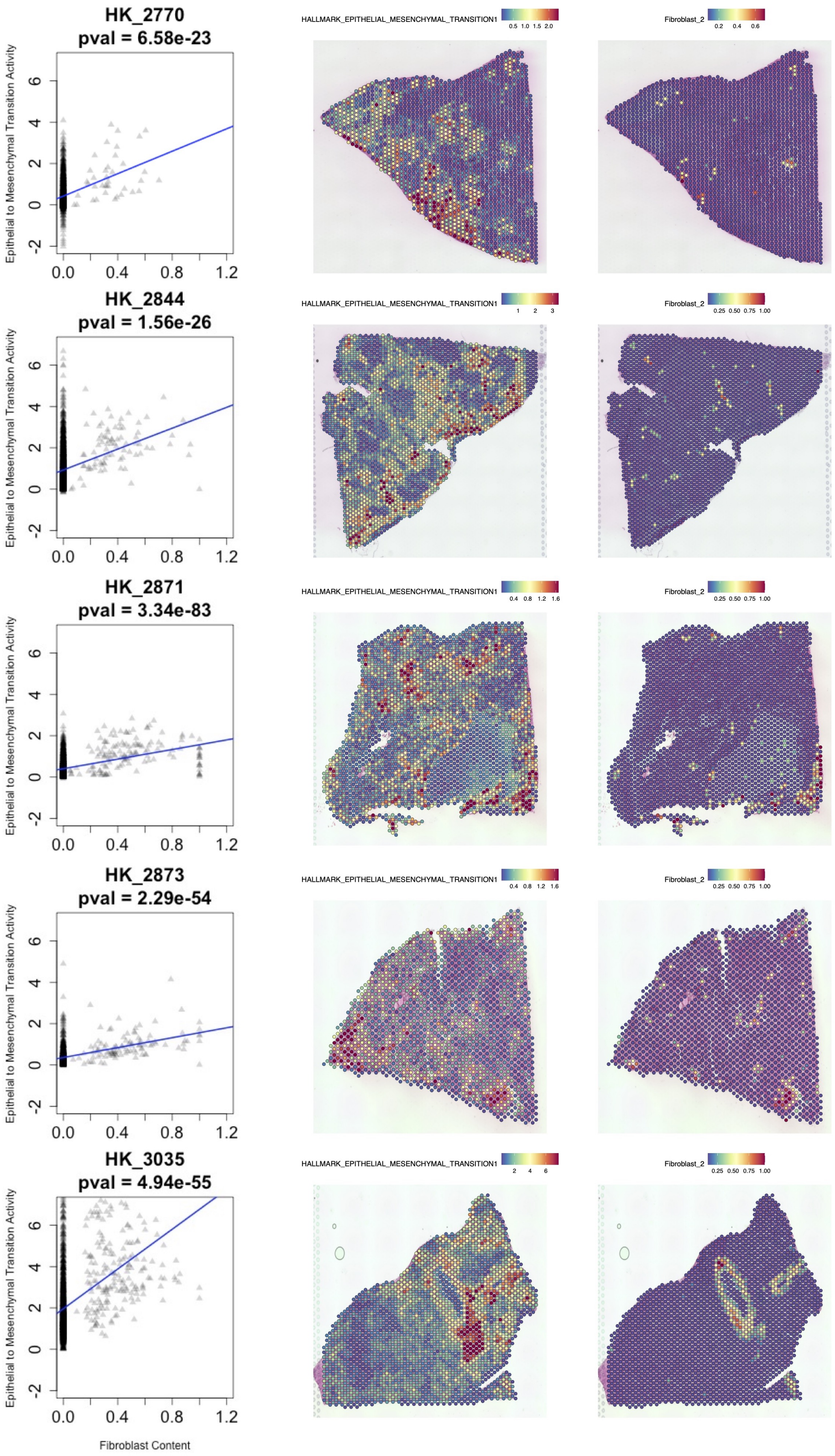


Fig. 14: Left: scatter plot of epithelial-mesenchymal transition (EMT ) pathway activity versus Fibroblast ratio. p-value is calculated from linear regression. Middle: epithelial-mesenchymal transition pathway activity. Right: fibroblast cells' abundance from deconvolution.

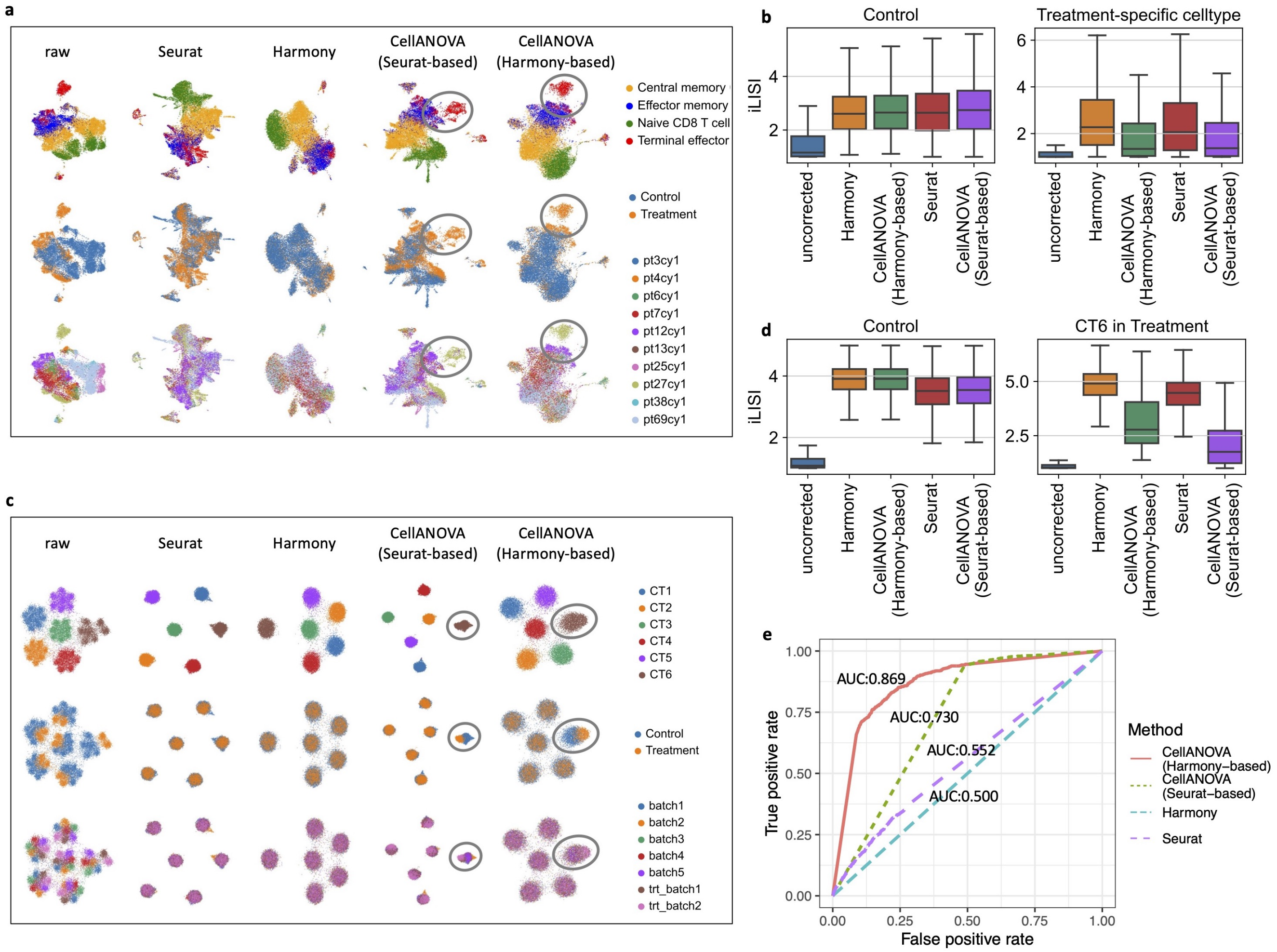


Fig. 15: (a) UMAP visualization of the "hold-one-celltype-out" experiment using immunotherapy data before and after batch correction, colored by cell type, condition, and batch. (b) Plot showing the LISI score of cells from the control pool (left) and cells belonging to treatment-specific cell type Terminal Effector (right) in the experiment in (a). (c) UMAP visualization of simulated data before and after batch correction, colored by cell type, condition, and batch. (d) Plot showing the LISI score of cells from the control pool (left) and cells belonging to treatment-specific cell type CT6 (right) in the simulation in (b). (e) ROC curves obtained from differential expression analysis between control and treatment groups, using the batch-corrected expressions of CT6 cells from different integration methods.

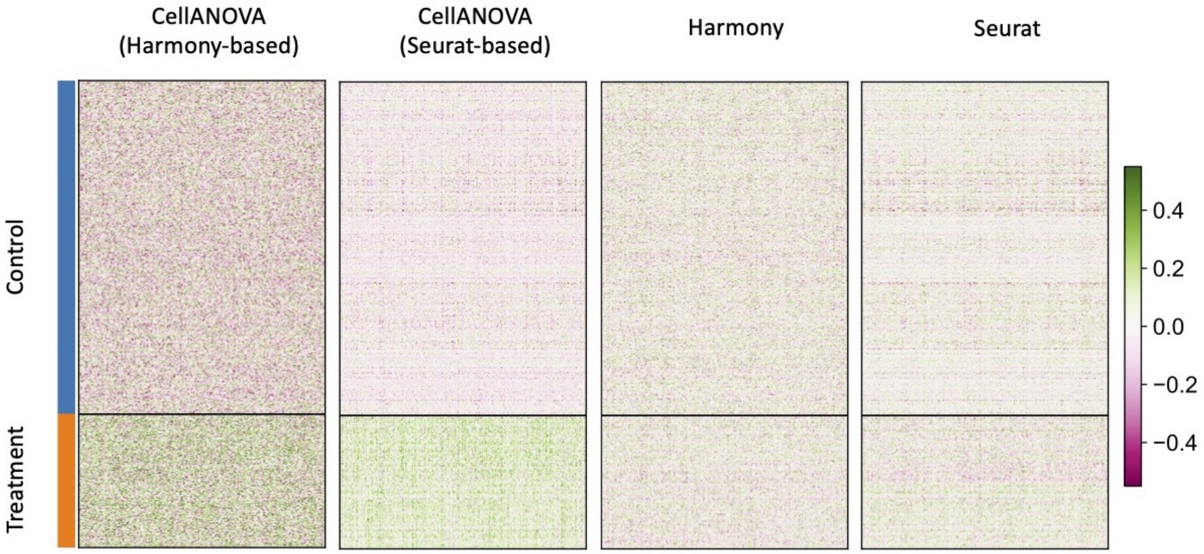


Fig. 16: Heatmaps of batch-corrected expressions of differentially expressed genes between control and treatment groups in the simulation study.

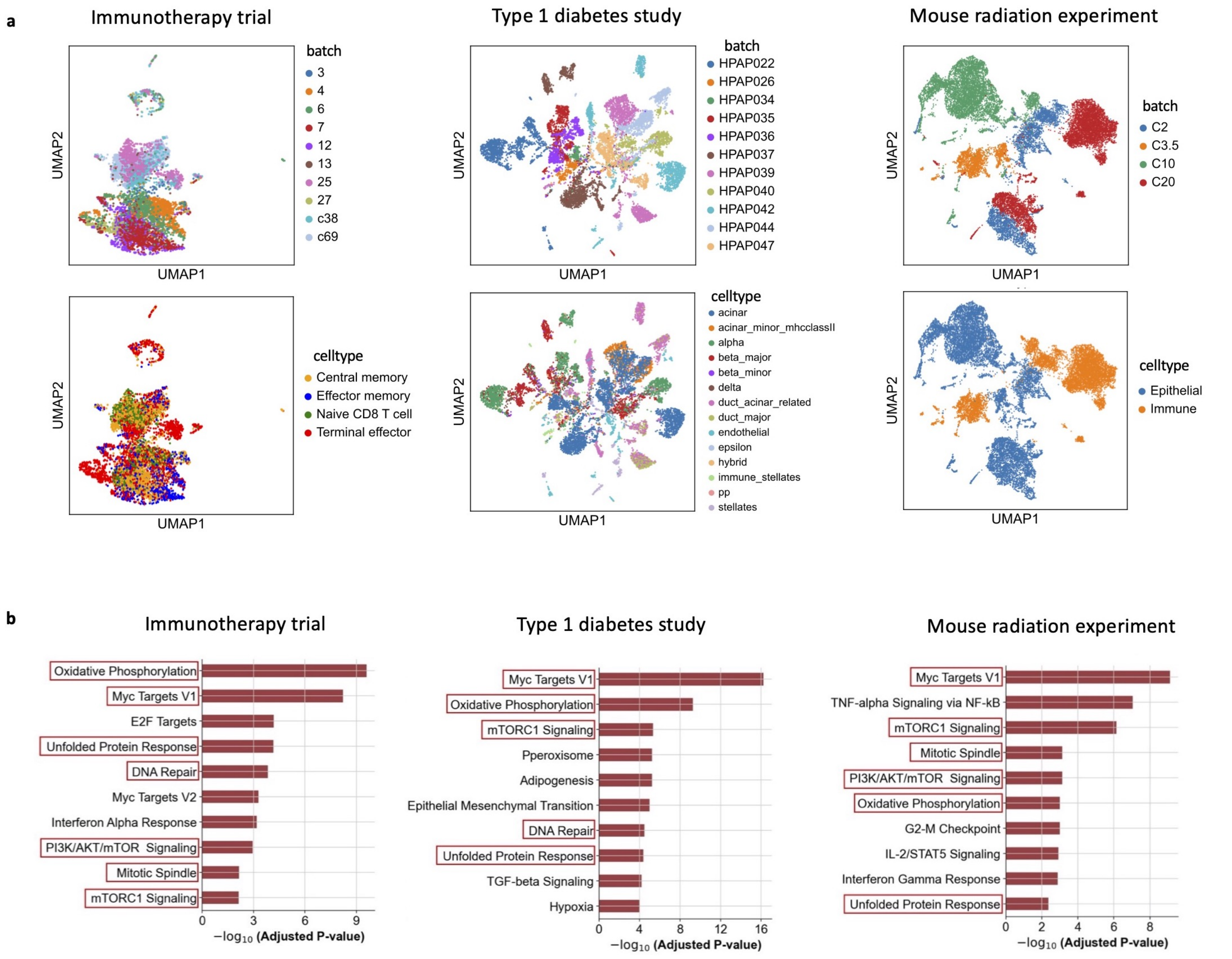


Fig. 17: (a) UMAP visualizations of batch effects estimated by Seurat-based CellANOVA on three datasets, colored by batch and cell type. (b) Top ten batch-affected pathways of each study based on batch-susceptibility score (BSS) with Seurat-based CellANOVA.

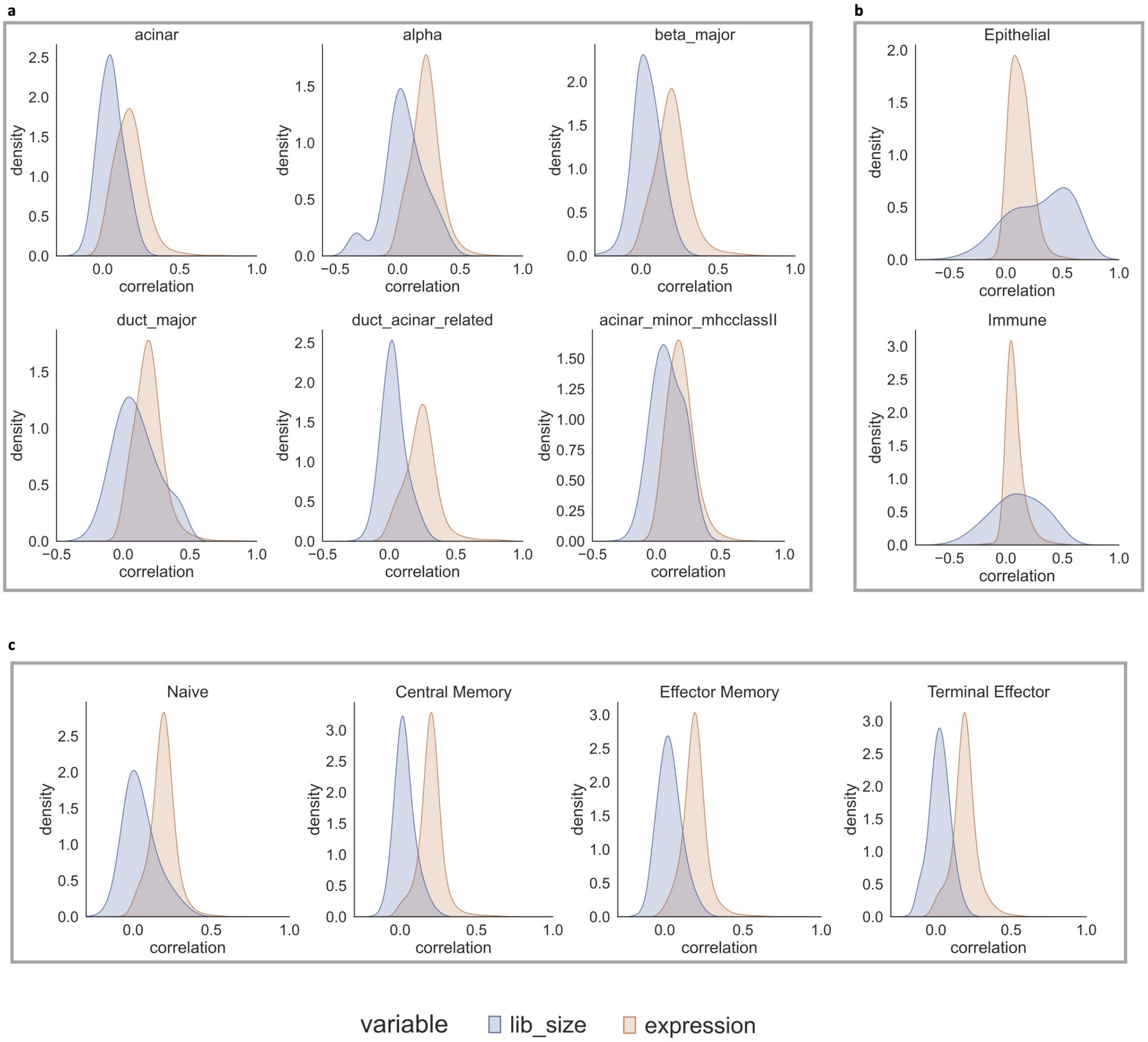


Fig. 18: Visualizations of the correlation between gene-specific batch effects, library size, and gene expression levels.
